# Increased levels of eIF2A inhibit translation by sequestering 40S ribosomal subunits

**DOI:** 10.1101/2022.11.18.517125

**Authors:** Daisy J. Grove, Daniel J. Levine, Michael G. Kearse

**Affiliations:** The Ohio State Biochemistry Program, Columbus, OH 43210 USA; Department of Biological Chemistry and Pharmacology, Columbus, OH 43210 USA; Center for RNA Biology, The Ohio State University, Columbus, OH 43210 USA

## Abstract

eIF2A was the first eukaryotic initiator tRNA carrier discovered but its exact function has remained enigmatic. Uncharacteristic of translation initiation factors, eIF2A is reported to be non-cytosolic in multiple human cancer cell lines. Attempts to study eIF2A mechanistically have been limited by the inability to achieve high yield of soluble recombinant protein. Here, we developed a purification paradigm that yields ∼360-fold and ∼6,000-fold more recombinant human eIF2A from *E. coli* and insect cells, respectively, than previous reports. Using a mammalian *in vitro* translation system, we found that increased levels of recombinant human eIF2A inhibit translation of multiple reporter mRNAs, including those that are translated by cognate and near-cognate start codons, and does so prior to start codon recognition. eIF2A also inhibited translation directed by all four types of cap-independent viral IRESs, including the CrPV IGR IRES that does not require initiation factors or initiator tRNA, suggesting excess eIF2A sequesters 40S subunits. Supplementation with additional 40S subunits prevented eIF2A-mediated inhibition and pull-down assays demonstrated direct binding between recombinant eIF2A and purified 40S subunits. These data support a model that eIF2A must be kept away from the translation machinery to avoid sequestering 40S ribosomal subunits.

## INTRODUCTION

Canonical mRNA translation initiation uses the ternary complex (TC)—comprised of the heterotrimeric eukaryotic initiation factor 2 (eIF2) bound to GTP and initiator tRNA (Met-tRNA_i_^Met^) —to deliver the initiator tRNA to the P site of the 40S ribosomal subunit. The TC along with eIF1, eIF1A, eIF3, eIF5, and the 40S subunit form the 43S pre-initiation complex (PIC) that is recruited to the 5’ m^7^G cap bound by the eIF4F complex (comprised of the cap-binding protein eIF4E, the scaffolding protein eIF4G, and the ATP-dependent RNA helicase eIF4A) (1). The PIC then scans 5ʹ-to-3ʹ in search of an AUG start codon (2,3). Once the initiator tRNA in the P site base pairs with the AUG start codon, eIF2 hydrolyzes GTP and releases the initiator tRNA, stimulating the dissociation of itself and most eIFs. The 60S subunit then joins, which is aided by eIF5B, to form the 80S ribosome (4). While eIF2 is the dominant Met-tRNA_i_^Met^ carrier in the cell, several other factors have been reported to bind and deliver Met-tRNA_i_^Met^ to the 40S subunit, namely eIF2A, eIF2D, and MCT-1•DENR (Multiple Copies in T-cell Lymphoma 1 and Density Regulated Protein complex) (5–7). Multiple reports suggest eIF2D and MCT-1•DENR function in translation re-initiation (6–8), but the exact role of eIF2A remains unknown (for an in-depth review on eIF2A see (9)).

eIF2A is a 65 kDa monomeric protein (non-homologous to eIF2) that was discovered over 50 years ago (initially named IF-M1) from the high salt-washed ribosome fraction of rabbit reticulocyte lysate (5,10). In 1975, Adams *et al*. used methionyl-puromycin synthesis assays to demonstrate that eIF2A was able to deliver Met-tRNA_i_^Met^ to the 40S subunit to form a functional 80S ribosome; however, eIF2A was much less efficient of a Met-tRNA_i_^Met^ carrier than eIF2 for endogenous mRNA (11). While eIF2A was first thought to be the functional ortholog of prokaryotic IF2 due to them both being monomeric and requiring a mRNA template to deliver Met-tRNA_i_^Met^, it soon became known that eIF2 (and not eIF2A) was the primary Met-tRNA_i_^Met^ carrier in eukaryotes. Intriguingly, endogenous eIF2A protein is as abundant than the eIF2 subunits (α, β, γ) in multiple human cell lines (12) (**Supplementary Figure S1A** and **Supplementary Table S1**) and has a similar reported binding affinity for Met-tRNA_i_^Met^ as eIF2, (12.4 nM and 15.0 nM, respectively) (13,14). However, little is known on a transcriptome wide level which mRNAs or open reading frames require eIF2A. CLIP-seq-based experiments to identify eIF2A-bound transcripts are lacking. To our knowledge, only a single published report has used ribosome profiling of eIF2A knockout (KO) cells, which shows a decrease in the ratio of upstream open reading frames (uORFs), many of which were non-AUG encoded, to main ORF translation; however, there was no global effect on translation upon KO (15).

Atypical of translation initiation factors, multiple reports have described eIF2A of being non-cytosolic in mammalian cells (14,16–18). For example, Kim *et al*. reported that eIF2A is primarily restricted to the nucleus during normal growth conditions but shuttles to the cytoplasm during cell stress and viral infection in Huh7 cells (14). Multiple reports have provided genetic evidence that eIF2A may selectively function in translation of specific mRNAs or ORFs, often at near-cognate start codons. Starck *et al*. demonstrated that eIF2A knockdown (KD) decreases the translation of the UUG-encoded uORF reporter in *Binding immunoglobin protein* (*BiP*) during cell stress (19). In an earlier report, Starck *et al*. concluded that Leu-tRNA^Leu^ can be used to initiate translation at CUG codons, which is not inhibited by NSC119893 that suppresses canonical eIF2-mediated initiation (20). KD of eIF2A decreased translation of CUG-encoded ORFs (20). These results led the authors to speculate that eIF2A can use Leu-tRNA^Leu^ for initiation (20); however, direct binding between eIF2A and Leu-tRNA^Leu^ was not confirmed. Liang *et al*. reported an isoform of *PTEN (PTEN-α)*, a gene commonly mutated in cancer, is translated from a CUG-encoded uORF and KD of eIF2A leads to decreased PTEN-α protein levels (21). Signal from repeat-associated non-AUG (RAN) translation reporters harboring myotonic dystrophy type 2 CCUG•CAGG repeats were reduced in eIF2A KO cells and increased when eIF2A was co-overexpressed in HEK293T cells (22). However, a steadfast role of eIF2A in all RAN translation is not clear as studies using different reporter designs have divergent conclusions of eIF2A depletion and RAN translation of *C9orf72* ALS-TFD GGGGCC repeats (23,24). eIF2A may also regulate IRES-mediated translation in yeast and human cells (14,25–27), although this is not entirely clear as conflicting results focusing on hepatitis C virus (HCV) and eIF2A have been reported (28). In yeast, eIF2A-null cells have unchanged polysome profiles compared to control strains; yet genetic and physical interactions with eIF5B and eIF4E supports that eIF2A functions in translation (25,29). eIF2A KO mice developed a metabolic syndrome and had decreased life spans by one year of age, suggesting eIF2A may have a role in aging (30).

Direct biochemical evidence supporting a role of eIF2A in translation initiation using AUG or near-cognate start codons is less evident since its initial discovery. Purifying eIF2A has historically been challenging and has slowed mechanistic interrogation of its function(s). The initial purification scheme for eIF2A from rabbit reticulocyte lysate (RRL) included eight steps and produced a seemingly homogenous final product but did not yield protein that was consistently active that could deliver labeled Met-tRNA_i_^Met^ to 40S subunits (5,10). Dmitriev *et al*. identified a co-purified factor (later identified and named eIF2D) that was also able to bind and deliver Met-tRNA_i_^Met^ to 40S subunits (6). Due to the high similarity in chromatographic properties between eIF2A and eIF2D, a new purification scheme was developed that selected for eIF2A but, for unknown reasons, led to inactive eIF2A that no longer had the ability to deliver initiator tRNA to the 40S subunit (6,10). Kim *et al*. reported the ability to produce 10 μg of recombinant human His6-eIF2A from 6 L of *E. coli* culture which had low nM affinity for initiator tRNA (14).

Here, we developed a robust method for production of recombinant human eIF2A from *E. coli* and insect cells with ∼360-fold and ∼6,000-fold higher yield, respectively. Titrating recombinant eIF2A into mammalian *in vitro* translation reactions inhibited translation of multiple reporter mRNAs, including those that are translated by cognate and near-cognate start codons, and reduced formation of 80S ribosomes and 48S initiation complexes, suggesting eIF2A inhibits translation prior to start codon recognition. Increased levels of eIF2A also inhibit translation directed by all four types of cap-independent viral IRESs, including those that do not require ribosomal scanning, initiation factors, or initiator tRNA, suggesting increased eIF2A sequesters the 40S subunit. Reactions supplemented with additional 40S subunits prevented inhibition and pull-down assays provide evidence of direct binding between recombinant eIF2A and purified 40S subunits. These data support a model that eIF2A must be kept away from the translation machinery to avoid sequestering 40S ribosomal subunits.

## MATERIALS AND METHODS

### Plasmids

The ORF for full-length human eIF2A (Ref seq RNA # NM_032025.5) was synthesized by Integrated DNA Technologies (IDT) and was cloned in pET His6 MBP TEV LIC cloning vector (1M), which was a gift from Scott Gradia (Addgene plasmid # 29656), through ligation-independent cloning (LIC) using Novagen’s LIC-qualified T4 DNA polymerase (Sigma # 70099-M) as described by Q3 Macrolab (http://qb3.berkeley.edu/macrolab/). The His6-tag was deleted from the N-terminus and inserted at the C-terminus. Mutations were achieved using the Q5 Site-Directed Mutagenesis Kit (NEB # E0552S). For recombinant expression and purification from insect cells, MBP-eIF2A-His6 was PCR amplified to incorporate a C-terminal FLAG tag and then subcloned into pFastBac1(Thermo Fisher # 10359016).

pcDNA3.1(+)/nLuc-3XFLAG was previously described (31). pcDNA3.1(+)/3XF-RLuc (with AUG, CUG, GUG, AAA start codon variants) was constructed by subcloning RLuc from pRL-SV40 (which was a kind gift from Aaron Goldstrohm; Promega # E2231) via PCR amplification to insert the N-terminal 3XFLAG tag. IRES-containing nLuc reporters were generated using an overlapping PCR method and cloned into pcDNA3.1(+) or pcDNA3-1D. The PV IRES template was pcDNA3 RLUC POLIRES FLUC and was a gift from Nahum Sonenberg (Addgene plasmid # 45642). The EMCV IRES and HCV IRES templates were kind gifts from Aaron Goldstrohm. pcDNA3.1-D/CrPV IGR IRES nLuc-3XFLAG was previously described (31) but was additionally modified to contain a strong hairpin (HP) upstream of the IRES element to block scanning pre-initiation complexes. All IRES reporters contained the same strong hairpin upstream of the IRES element (which is noted in the complete reporter sequence in the **Supplementary Data**). Hairpin insertion (for IRES reporters) and all mutations were introduced using the Q5 Site-Directed Mutagenesis Kit (NEB # E0554S). We have previously published the use of the four HP-containing viral IRES nLuc reporters (32).

All plasmids were propagated in TOP10 *E. coli* (Thermo Fisher # C404006), purified using the PureYield Plasmid Miniprep or Midiprep Systems (Promega # A1222 and A2495), and validated by Sanger sequencing at The Ohio State University Comprehensive Cancer Center Genomics Shared Resource (OSUCCC GSR). Nucleotide sequences of all reporters and full-length recombinant proteins are provided in the **Supplementary Data**.

### Recombinant protein expression and purification

*E. coli* derived recombinant His6-MBP and MBP-eIF2A-His6 were produced in Rosetta 2(DE3) *E. coli* (Sigma # 71397-4) using MagicMedia *E. coli* Expression Medium (Thermo Fisher # K6803) supplemented with 50 µg/mL kanamycin and 35 µg/mL chloramphenicol for auto-induction. A 5 mL starter culture in LB media supplemented with 50 µg/mL kanamycin, 35 µg/mL chloramphenicol, and 1% glucose (w/v) was inoculated with a single colony and grown overnight at 37°C, 250 rpm. 1 mL of fresh overnight starter culture was then used to inoculate 50 mL of room temperature MagicMedia with 50 µg/mL kanamycin and 35 µg/mL chloramphenicol, and incubated for 72 hrs at 18°C, 160 rpm in a 250 mL baffled flask. After auto-induction, cultures were pelleted and stored at −20°C. Recombinant proteins were purified using a dual affinity approach, first using the C-terminal His6-tag, then the N-terminal MBP-tag. Cell pellets were resuspended and lysed with BugBuster Master Mix (Sigma # 71456) using the recommended 5 mL per 1 g wet cell pellet ratio for 10 min at room temperature with gentle end-over-end rotation (10-15 rpm). Lysates were placed on ice and kept cold moving forward. Lysates were cleared by centrifugation for 20 min at 18,000 rcf in a chilled centrifuge (4°C) and then incubated with HisPur Cobalt Resin (Thermo Fisher # 89965) in a Peirce centrifugation column (Thermo Fisher # 89897) for 30 min at 4°C with gentle end-over-end rotation. Columns were centrifuged in a pre-chilled (4°C) Eppendorf 5810R for 2 min at 700 rcf to eliminate the flow through and then were washed 5X with two resin bed volumes of ice-cold Cobalt IMAC Wash Buffer (50 mM Na_3_PO_4_, 300 mM NaCl, 10 mM imidazole; pH 7.4) in a pre-chilled (4°C) Eppendorf 5810R for 2 min at 700 rcf. His-tagged proteins were then eluted in a single elution step with two resin bed volumes of ice-cold Cobalt IMAC Elution Buffer (50 mM Na_3_PO_4_, 300 mM NaCl, 150 mM imidazole; pH 7.4) by gravity flow. Eluates were then incubated with Amylose Resin (NEB # E8021) in a centrifugation column for 2 hrs at 4°C with gentle end-over-end rotation. Columns were washed 5X with at least two resin bed volumes of ice-cold MBP Wash Buffer (20 mM Tris-HCl, 200 mM NaCl, 1 mM EDTA; pH 7.4) by gravity flow. MBP-tagged proteins were then eluted by a single elution step with two resin bed volumes of ice-cold MBP Elution Buffer (20 mM Tris-HCl, 200 mM NaCl, 1 mM EDTA, 10 mM maltose; pH 7.4) by gravity flow. Recombinant proteins were then desalted and buffer exchanged into Protein Storage Buffer (25 mM Tris-HCl, 125 mM KCl, 10% glycerol; pH 7.4) using a 7K MWCO Zeba Spin Desalting Column (Thermo Fisher # 89892) and, if needed, concentrated using a 10K MWCO Amicon Ultra-4 Filter Unit (EMD Millipore # UFC803024). Recombinant protein concentration was determined by Pierce Detergent Compatible Bradford Assay Kit (Thermo Fisher # 23246) with BSA standards diluted in Protein Storage Buffer before aliquoting in single use volumes, snap freezing in liquid nitrogen, and storage at −80°C.

Insect cell derived recombinant His6-mEGFP-FLAG and MBP-eIF2A-His6-FLAG was expressed in ExpiSf9 cells using the ExpiSf Expression System Starter Kit (Thermo Fisher # A38841) following the manufacturer’s protocol. Bacmids were produced by transforming MAX Efficiency DH10Bac Competent Cells (Thermo Fisher # 10361012) and selecting for integrated transformants (white colonies) on LB Agar supplemented with 50 μg/mL kanamycin, 7 μg/mL gentamicin, 10 μg/mL tetracycline, 300 μg/mL Bluo-gal, 40 μg/mL IPTG. Single integrated transformants were then re-streaked on selective LB Agar and then used to inoculate 100 mL of LB supplemented with 50 μg/mL kanamycin, 7 μg/mL gentamicin, 10 μg/mL tetracycline with overnight incubation at 37°C, 250 rpm. Bacmids were isolated from 30 mL of culture using PureLink HiPure Plasmid Midiprep Kit (Thermo Fisher # K210004). 25 mL of ExpiSf9 cells at 2.5 x 10^6^ cells/mL in ExpiSf CD medium in a 125 mL non-baffled vented PETG flask (Thermo Fisher # 4115-0125) were transfected with 12.5 µg of bacmid using 30 µL ExpiFectamine Sf Transfection Reagent in 1 mL Opti-MEM I Reduced Serum Media and incubated at 27°C, 195 rpm. P0 baculovirus stocks were collected after 5 days and stored at 4°C for less than a week before long-term storage at −80°C. For protein expression, 240 mL of ExpiSf9 cells at 5 x 10^6^ cells/mL in ExpiSf CD medium in a 1L non-baffled vented PETG flask (Thermo Fisher # 4115-1000) were treated with 800 µL ExpiSf Enhancer and incubated at 27°C, 195 rpm for 22 hr. 3 mL of P0 baculovirus stock was then added and allowed to incubate for 72 hrs at 27°C, 195 rpm. Cells were then harvested (50 mL culture pellets), snap frozen in liquid nitrogen, and stored at −80°C. A single cell pellet from 50 mL of culture was then lysed in 16 mL (4 pellet volumes) of ice-cold Lysis Buffer (25 mM Tris, 300 mM KCl, 10% (v/v) glycerol, 0.5% (v/v) Igepal CA-630, protease inhibitor EDTA-free (Thermo Fisher # A32955), phosphatase inhibitor cocktail (Thermo Fisher # A32957); pH 7.5) with gentle end-over-end rotation for 15 min at room temperature. Lysates were cleared by centrifugation at 214,743 rcf at 4°C in S55A rotor using a Sorvall Discovery M120 SE Micro-Ultracentrifuge and added to 1 mL of Pierce Anti-DYKDDDDK Affinity Resin (Thermo Fisher # A36803) in Peirce centrifugation column (Thermo Fisher # 89897) for 2 hrs at 4°C with gentle end-over-end rotation. Columns were washed 4X with at least two resin bed volumes of ice-cold Lysis Buffer and once with room temperature Elution Buffer without Peptide (25 mM Tris, 125 mM KCl, 10% (v/v) glycerol; pH 7.5) by gravity flow. FLAG-tagged proteins were then eluted with room temperature Elution Buffer with 2.5 mg/mL 3XFLAG Peptide (Thermo Fisher # A36806) for 15 min at room temperature. Eluates were placed on ice and then incubated with Amylose Resin (NEB # E8021) in a centrifugation column for 2 hrs at 4°C with gentle end-over-end rotation. Columns were washed 5X with at least two bed volumes of ice-cold MBP Wash Buffer (20 mM Tris-HCl, 200 mM NaCl, 1 mM EDTA; pH 7.4) by gravity flow. MBP-tagged proteins were then eluted by a single elution step with two resin bed volumes of ice-cold MBP Elution Buffer (20 mM Tris-HCl, 200 mM NaCl, 1 mM EDTA, 10 mM maltose; pH 7.4) by gravity flow. Recombinant proteins were then desalted and buffer exchanged into Protein Storage Buffer (25 mM Tris-HCl, 125 mM KCl, 10% glycerol; pH 7.4) using a 7K MWCO Zeba Spin Desalting Column (Thermo Fisher # 89892). Recombinant protein concentration was determined by Pierce Detergent Compatible Bradford Assay Kit (Thermo Fisher # 23246) with BSA standards diluted in Protein Storage Buffer before aliquoting in single use volumes, snap freezing in liquid nitrogen, and storage at −80°C.

Insect cell derived recombinant tag-less eIF2A was expressed in ExpiSf9 cells using the ExpiSf Expression System Starter Kit (Thermo Fisher # A38841) following the manufacturer’s protocol and adapting the IMPACT (Intein Mediated Purification with an Affinity Chitin-binding Tag) System (NEB # 6901S). The human eIF2A coding sequence was cloned into the C-terminal Mxe GyrA Intein-chitin-binding domains (CBD) expression vector pTXB1 and then subcloned into pFastBac1. Bacmid and P0 baculovirus stock generation, along with ExpiSf9 transduction and expression was performed as described above. A single cell pellet from 50 mL of culture was then lysed in 16 mL (4 pellet volumes) of ice-cold Lysis Buffer (20 mM Tris, 500 mM KCl, 10% (v/v) glycerol, 0.5% (v/v) Igepal CA-630, protease inhibitor EDTA-free (Thermo Fisher # A32955), phosphatase inhibitor cocktail (Thermo Fisher # A32957); pH 8.0) with 7 kU Pierce Universal Nuclease for Cell Lysis (Thermo Fisher # 88702) and gentle end-over-end rotation for 15 min at room temperature. Lysates were cleared by centrifugation at 214,743 rcf at 4°C in S55A rotor using a Sorvall Discovery M120 SE Micro-Ultracentrifuge and added to 1 mL of Chitin Resin (NEB # S6651S) in a Peirce centrifugation column (Thermo Fisher # 89897) for 2 hrs at 4°C with gentle end-over-end rotation. Columns were washed 10X with 1 column volume of Chitin Resin Wash Buffer (20 mM Tris-HCl, 500 mM KCl, 10% (v/v) glycerol, 0.5% (v/v) Igepal CA-630; pH 8.0). Three resin bed volumes of Cleavage Buffer (20 mM Tris-HCl, 500 mM KCl, 50 mM DTT; pH 8.0) was dripped through via gravity flow once until a small amount of buffer remained above the resin. The flow was stopped by capping the column. DTT-induced on column intein cleavage proceeded at 22°C for 16 hours with no shaking or rotation. The column was then dripped by gravity flow into a pre-chilled tube and labeled as eluate 1. Recombinant proteins were eluted 6X with two resin bed volumes of Wash Buffer by gravity flow with each eluate being collected in a separate pre-chilled tube (labeled as eluates 2-7). Eluates were then analyzed by SDS-PAGE and Coomassie staining. Eluates 1 and 2 typically contained the most abundant and pure cleaved protein, which were then pooled and concentrated using a 10K MWCO Amicon Ultra-4 Filter Unit (EMD Millipore # UFC803024). Recombinant proteins were then desalted into Protein Storage Buffer (25 mM Tris-HCl, 125 mM KCl, 10% glycerol; pH 7.4) using a 7K MWCO Zeba Spin Desalting Column (Thermo Fisher # 89892). Recombinant protein concentration was determined by Pierce Detergent Compatible Bradford Assay Kit (Thermo Fisher # 23246) with BSA standards diluted in Protein Storage Buffer before aliquoting in single use volumes, snap freezing in liquid nitrogen, and storage at −80°C.

Human cell derived recombinant eIF2A-FLAG was expressed in HEK293T cells obtained commercially (OriGene # TP304303) and buffer exchanged into Protein Storage Buffer (25 mM Tris-HCl, 125 mM KCl, 10% glycerol; pH 7.4) using a 7K MWCO Zeba Spin Desalting Column (Thermo Fisher # 89892). Recombinant protein concentration was determined by Pierce Detergent Compatible Bradford Assay Kit (Thermo Fisher # 23246) with BSA standards diluted in Protein Storage Buffer before aliquoting in single use volumes, snap freezing in liquid nitrogen, and storage at −80°C.

### *In vitro* transcription

nLuc-3XFLAG, 3XFLAG-RLuc, β-globin 5ʹ UTR-nLuc-3XFLAG, and ATF4 5ʹ UTR-nLuc-3XFLAG reporter plasmids were linearized with PspOMI. PV IRES nLuc-3XFLAG, EMCV IRES nLuc-3XFLAG, HCV IRES nLuc-3XFLAG, and CrPV IGR IRES nLuc-3XFLAG reporter plasmids were linearized with XbaI. Digested plasmids were purified using the DNA Clean and Conentrator-25 (Zymo Research # 11-305C). 0.5 μg of linearized plasmid was used as template in a 10 µL reaction using the HiScribe T7 High Yield RNA Synthesis Kit (NEB # E2040S) with an 8:1 ratio of cap analog to GTP, producing ∼90% capped RNA, for 2 h at 30°C. Template DNA was digested with the addition of 1 μL RNase-free DNaseI (NEB # M0303S; enzyme stock at 2,000 U/mL) for 15 min at 37°C. mRNAs were subsequently polyadenylated using *E. coli* poly(A) polymerase (NEB # M0276S) with the addition of 5 μL 10X buffer, 5 μL 10 mM ATP, 1 μL *E. coli* poly(A) polymerase (enzyme stock at 5,000 U/mL), and 28 μL RNase-free water for 1 hr at 37°C. mRNAs were purified using RNA Clean and Concentrator-25 (Zymo Research # 11-353B), eluted in 75 µL RNase-free water, aliquoted in single-use volumes, and stored at −80°C. nLuc-3XFLAG, 3XFLAG-RLuc, β-globin 5ʹ UTR-nLuc-3XFLAG, and ATF4 5ʹ UTR-nLuc-3XFLAG mRNAs were co-transcriptionally capped with the 3’-O-Me-m7G(5ʹ)ppp(5ʹ)G RNA Cap Structure Analog (NEB # S1411S). All viral IRES nLuc-3XFLAG mRNAs were co-transcriptionally capped with the A(5ʹ)ppp(5ʹ)G RNA Cap Structure Analog (NEB # S1406S).

### *In vitro* translation and luciferase assays

10 μL *in vitro* nLuc mRNA translation reactions were performed in the dynamic linear range using 3 nM mRNA (31) in the Flexi Rabbit Reticulocyte Lysate (RRL) System (Promega # L4540) with final concentrations of reagents at 20% RRL, 10 nM amino acid mix minus Leucine, 10 nM amino acid mix minus Methionine, 100 mM KCl, 0.5 mM MgOAc, 8 U murine RNase inhibitor (NEB # M0314L), and 0-3.4 μM recombinant protein. The final KCl concentration was kept constant and was accounted for in the Protein Storage Buffer. Reactions were pre-incubated with recombinant protein for 10 min on ice before the addition of mRNA, then incubated for 30 min at 30°C and terminated by incubation on ice. nLuc luciferase signal was measured by mixing 25 μL of Nano-Glo Luciferase Assay System (prepared 1:50 as recommended; Promega # N1120) with 25 μL of diluted reactions (diluted 1:5 in Glo Lysis Buffer; Promega # E2661) and incubated at room temperature for 5 min, and then read on a Promega GloMax Discover multimode plate reader. RLuc mRNA translation reactions were performed identically as described above but were incubated at 30°C for 90 min (this allowed the weaker RLuc enzyme to give signal above background for the start codon mutants) and diluted with 25 μL Glo Lysis Buffer. 25 μL diluted lysate was mixed with an equal volume of *Renilla*-Glo Luciferase Assay System (prepared 1:100 as recommended; Promega # E2710) for 10 min at room temperature.

### Western blotting

10 µL translation reactions were performed as described above, then mixed with 40 µL of 2X reducing LDS sample buffer (Bio-Rad # 1610747) and heated at 70°C for 15 min. 15 µL was then separated by standard Tris-Glycine SDS-PAGE (Thermo Fisher # XP04200BOX) and transferred on to 0.2 µm PVDF membrane (Thermo Fisher # 88520). Membranes were then blocked with 5% (w/v) non-fat dry milk in TBST (1X Tris-buffered saline with 0.1% (v/v) Tween 20) for 30 min at room temperature before overnight incubation with primary antibodies in TBST at 4°C. After three 10 min washes with TBST, membranes were incubated with HRP-conjugated secondary antibody in TBST for 1 hr at room temperature and then washed again with three 10 min washes with TBST. Chemiluminescence was performed with SuperSignal West Pico PLUS (Thermo Fisher # 34577) imaged using an Azure Sapphire Biomolecular Imager. Rabbit anti-GAPDH was used at 1:1,000 (Cell Signaling # 5174S). Rabbit anti-RPS3 (Bethyl # A303-840A) was used at 1:1,000. Rabbit anti-RPS6 (Cell Signaling # 2217S) was used at 1:1,000. Rabbit anti-RPL7 was used at 1:1,000 (Abcam # ab72550). Mouse anti-FLAG was used at 1:1,000 (Sigma # F1804). HRP-conjugated goat anti-rabbit IgG (H+L) (Thermo Fisher # 31460) and HRP-conjugated goat anti-mouse (Thermo Fisher # 31430) were both used at 1:10,000 for all blots, with the only exceptions being a 1:20,000 dilution for anti-RPS3 blots and a 1:30,000 dilution for anti-RPL7 blots.

### Sucrose gradient ultracentrifugation, RNA extraction, and RT-qPCR

*In vitro* translation reactions were programmed as described above with 3 nM nLuc mRNA (final) except were scaled up to 100 μL. Reactions were spiked with 50 µM lactimidomycin (5 mM stock in DMSO; Millipore # 5.06291.001) or 5 mM GMPPNP (stock at 100 mM in 100 mM Tris-HCl, pH 7.7; Sigma # G0635-5MG). Reactions were pre-incubated with inhibitors without recombinant protein or mRNA for 10 min at 30°C, then placed on ice. The indicated recombinant proteins were added and reactions were incubated for 10 min on ice. mRNA was added and 48S initiation complexes and 80S ribosomes were allowed to form for 10 min at 30°C, then returned to ice. Reactions with GMPPNP were supplemented with additional Mg^2+^ (1 mM final) to balance the cation sequestered by the added GMPPNP (this was optimized by determining 48S abundance with control reactions with native RRL in sucrose gradients). As a negative control, an mRNA only sample of 100 µL 3 nM reporter mRNA in RNase-free water was used. Reactions were then diluted with 100 µL (equal volume) of ice-cold 2X Polysome Dilution Buffer (40 mM Tris-HCl, 280 mM KCl, 10 mM MgCl_2_, 200 µg/mL cycloheximide, 2 mM DTT; pH 7.5) and layered on top of a linear 5-30% (w/v) buffered sucrose gradient (10 mM Tris-HCl, 140 mM KCl, 10 mM MgCl_2_, 100 μg/mL cycloheximide, 1 mM DTT; pH 7.5) in a 14 mm × 89 mm thin-wall Ultra-Clear tube (Beckman # 344059) that was formed using a Biocomp Gradient Master. Gradients were centrifuged at 35K rpm for 3 hrs at 4°C in a SW-41Ti rotor (Beckman) with maximum acceleration and no brake using a Beckman Optima L-90 Ultracentrifuge. Gradients were subsequently fractionated into 0.5 mL volumes using a Biocomp piston fractionator with a TRIAX flow cell (Biocomp) recording a continuous A_260 nm_ trace. Total RNA was extracted from 400 μL of each fraction (spiked with 0.2 ng exogenous control FFLuc mRNA; Promega # L4561) by adding 600 μL TRIzol (Thermo Fisher # 15596018) and following the manufacturer’s protocol. Glycogen (Thermo Fisher # R0561) was added at the isopropanol precipitation step. The resulting RNA pellet was resuspended in 30 μL nuclease-free water. 16 μL of extracted RNA was converted to cDNA using iScript Reverse Transcription Supermix for RT-qPCR (Bio-Rad # 1708841). cDNA reactions were then diluted 10-fold with nuclease-free water and stored at −20°C or used immediately. RT-qPCR was performed in 15 μL reactions using iTaq Universal SYBR Green Supermix (Bio-Rad # 1725124) in a Bio-Rad CFX Connect Real-Time PCR Detection System with 1.5 μL diluted cDNA and 250 nM (final concentration) primers. For each fraction, nLuc reporter mRNA abundance was normalized to the spiked-in control FFLuc mRNA using the Bio-Rad CFX Maestro software (ΔΔCt method). Abundance of total signal in each fraction was calculated using *Q_n_ = 2^ΔΔCt^*and *P = 100 × Q_n_/Q_total_* as previously described (33). Primers for RT-qPCR can be found in **Supplementary Table S2**.

### 40S ribosomal subunit and 80S ribosome purification

250 mL of native rabbit reticulocyte lysate (Green Hectares) was thawed overnight at 4°C, transferred to 25 x 89 mm Ultra-Clear thin-wall tubes (Beckman # 344058) (typically 38 mL per tube) and centrifuged at 28,000 rpm (140,992.2 rcf) for 4 hours in a SW28 rotor using a Beckman Coulter Optima L-90K Ultracentrifuge. The supernatant was aspirated and each crude ribosome pellet was resuspended in 500 μL ice-cold Buffer A (20 mM HEPES, 500 mM KCl, 1 mM MgCl_2_, 0.25 M sucrose; pH 7.5) by gentle orbital shaking overnight at 4°C in the dark and then gentle manual pipetting. Resuspensions were combined in a 1.7 mL microcentrifuge tube, gently mixed using end-over-end rotation (12 rpm) for 30 min at 4°C, and centrifuged at 18,000 rcf for 10 min at 4°C. Supernatants were pooled and puromycin was added to a final concentration of 5 mM, incubated 15 min on ice, followed by gentle mixing for 15 min at 37°C to separate ribosomal subunits, and then snap frozen in liquid nitrogen and stored at −80°C. Puromycin-treated crude ribosomes were thawed on ice, diluted 1:4 with ice-cold Buffer B (20 mM HEPES, 500 mM KCl, 1 mM MgCl_2_, 2 mM DTT; pH 7.5) and 400 μL was loaded onto a 5-30% (w/v) buffered sucrose gradient (in Buffer B) in a 14 mm × 89 mm thin-wall Ultra-Clear tube (Beckman # 344059) that was formed using a Biocomp Gradient Master. Six gradients were centrifuged at 35K rpm for 3 hrs at 4°C in a Beckman SW-41Ti rotor with maximum acceleration and no brake using a Beckman Optima L-90 Ultracentrifuge. Gradients were subsequently fractionated into 0.5 mL volumes using a Biocomp piston fractionator with a TRIAX flow cell (Biocomp) recording a continuous A_260 nm_ trace. Fractions that contained the lighter edge of the 40S peak (i.e., only the early fractions that contained the 40S peak to avoid eIF3-bound 40S subunits) were pooled and pelleted in a 11 x 34 mm thin-wall polycarbonate tube (Thermo Fisher # 45315) at 55,000 rpm for 18 hrs at 4°C in a Sorvall S55-S rotor using a Sorvall Discovery M120 SE Micro-Ultracentrifuge. The supernatant was removed and each pellet was resuspended in 25 µL ice-cold Buffer C (20 mM HEPES, 100 mM KCl, 2.5 mM MgCl_2_, 0.25 M sucrose, 2 mM DTT; pH 7.5) and pooled. A_260_ values were measured using a Nanodrop spectrophotometer and concentration was calculated using 1 A_260_ unit = 65 pmol/mL (34). Purified 40S subunits were aliquoted into 5 μL volumes, flash frozen in liquid nitrogen, and stored at −80°C.

80S ribosomes were purified from the high salt-wash ribosome fraction from RRL as above, except samples were not treated with puromycin and were not incubated at 37°C. Fractions in the 5-30% (v/v) buffered sucrose gradient (in Buffer B) corresponding to the 80S ribosome were pooled and pelleted overnight as described above. A_260_ values were measured using a Nanodrop spectrophotometer and concentration was calculated using 1 A_260_ unit = 20 pmol/mL (35). Purified 80S ribosomes were aliquoted into 5 μL volumes, flash frozen in liquid nitrogen, and stored at −80°C.

### His6 pulldown binding experiments

In 20 µL, 3 µg recombinant His6-MBP or MBP-eIF2A-His6 was mixed with 20% RRL in 100 mM KCl, 10 nM amino acid mix minus methionine, 10 nM amino acid mix minus leucine, and 0.5 mM MgOAc (same as *in vitro* translation above) or with 0.27 μM purified 40S ribosomal subunits in 40S Binding Buffer (14) (final of 50 mM HEPES, 100 mM KCl, 5 mM MgCl_2_,; pH 7.5) and were incubated at 25°C for 10 min. Indicated samples were then crosslinked with 0.25% (v/v) formaldehyde (Sigma # F79-500) for 30 min at 25°C and quenched with 20 µL 100 mM Tris-HCl, pH 7.5. Samples were diluted 1:10 in ice-cold Wash Buffer (20 mM Tris-HCl, 140 mM KCl, 10 mM MgCl_2_, 0.1% (v/v) Triton X-100, 10 mM imidazole; pH 7.5) and added to 20 μL HisPur Ni-NTA Magnetic Beads (Thermo Fisher # 888831) for 30 min at 4°C with end-over-end rotation. For 40S ribosomal subunit and 80S ribosome binding experiments, HisPur Ni-NTA Magnetic Beads were blocked with 1 μg/μL BSA (Invitrogen # AM2616) and 2 μg/mL yeast tRNA (Thermo Fisher # AM7119) in Wash Buffer for 1 hr at 4°C with end-over-end rotation. Beads were then washed 5X with 400 µL ice-cold Wash Buffer. Bound proteins were then eluted with 200 µL 2X reducing LDS sample buffer (BioRad # 1610747) and heating for 15 min at 70°C. 20 µL of eluate was analyzed by SDS-PAGE and Western blotting as described above.

## RESULTS

### Recombinant eIF2A inhibits AUG- and near-cognate-initiated translation *in vitro*

Native eIF2A can be purified from rabbit reticulocyte lysate (often contaminated with eIF2D) (5,6,10). eIF2A can also be expressed recombinantly in *E. coli* and purified to high purity but requires large volumes of culture for little yield (i.e., 6 L of culture yields 10 μg soluble protein) (14). To overcome these barriers, we first adapted an autoinduction expression system using Rosetta 2(DE3) *E. coli* cells and dual-tagged human eIF2A to purify only full-length MBP-eIF2A-His6 protein (**Figure 1A**). By comparison, a single 50 mL autoinduction culture yielded 31.2 µg (an ∼ 360-fold increase in yield/mL culture). We consistently observed a sub-stoichiometric amount of *E. coli* GroEL chaperone (identified by mass spectrometry) co-eluting with recombinant eIF2A. Compared to the His6-MBP tag alone that accumulates to very high levels, MBP-eIF2A-His6 did not overexpress well. Switching MBP for GST or SUMO did not increase expression; however, fusing NusA to eIF2A allowed very large amounts of insoluble NusA-eIF2A-His6 to accumulate (data not shown).

**Figure 1.**
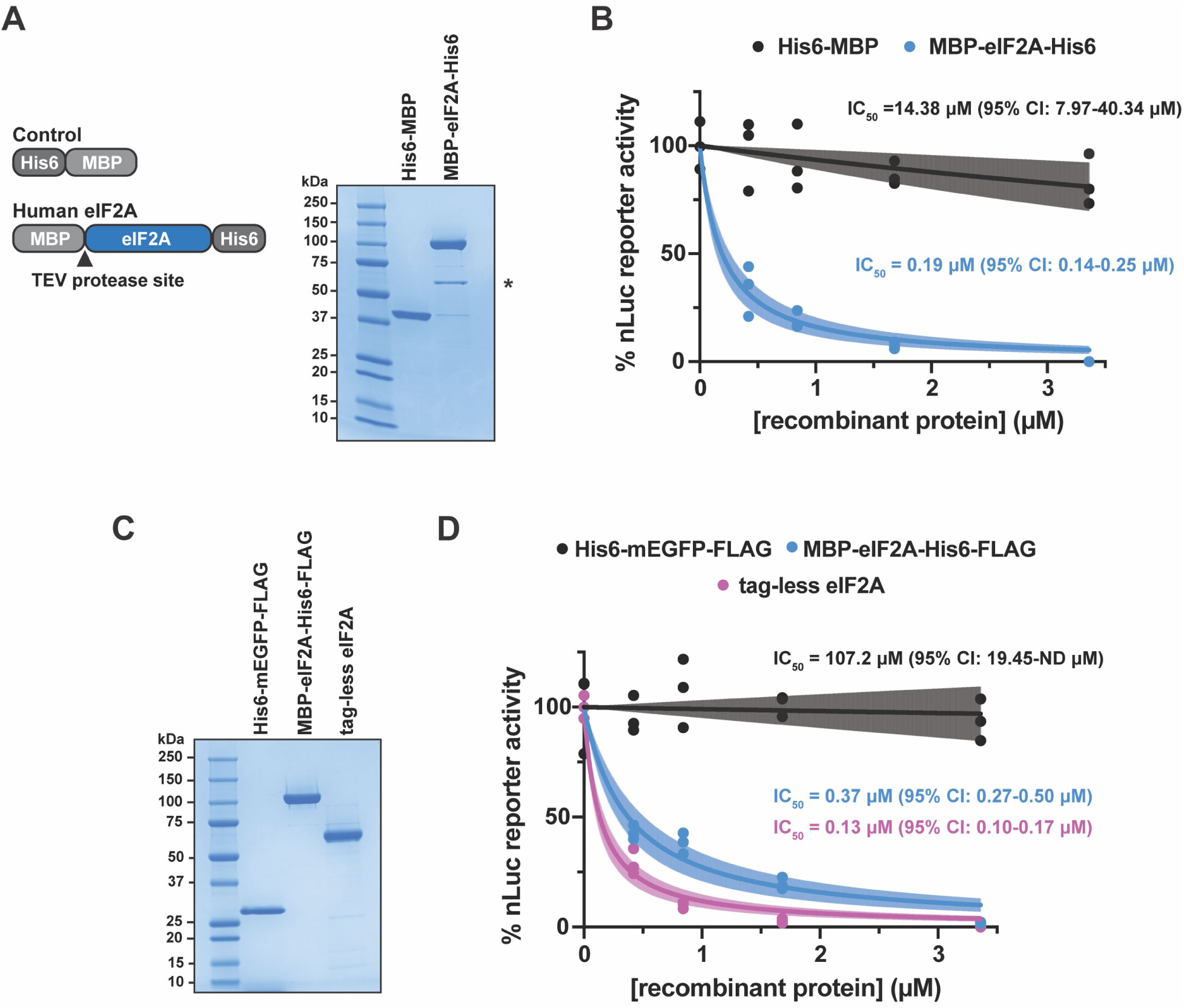
Recombinant eIF2A inhibits translation *in vitro*. A) Schematic of His6-MBP and MBP-eIF2A-His6 (left). SDS-PAGE and Coomassie stain of recombinant His6-MBP and MBP-eIF2A-His6 (right). 2 μg of protein was loaded. * = co-purified bacterial GroEL chaperone. B) *In vitro* translation of nLuc mRNA with a titration (0-3.4 μM) of the indicated recombinant proteins. IC_50_ values were determined for His6-MBP (14.38 μM with 7.97-40.34 μM 95% CI) and MBP-eIF2A-His6 (0.19 μM with 0.14-0.25 μM 95% CI). n=3 biological replicates. A non-linear regression was used to calculate the IC_50_ and is shown as the line with the 95% confidence interval (CI) included as a watermark. C) SDS-PAGE and Coomassie stain of insect cell derived His6-mEGFP-FLAG, MBP-eIF2A-His6-FLAG, and tag-less eIF2A. 2 μg of protein was loaded. D) *In vitro* translation of nLuc mRNA with a titration (0-3.4 μM) of insect cell-synthesized recombinant protein. IC_50_ values were determined for His6-mEGFP-FLAG (107.2 μM with 19.45-ND μM 95% CI), MBP-eIF2A-His6-FLAG (0.37 μM with 0.27-0.50 μM 95% CI), and tag-less eIF2A (0.13 μM with 0.10-0.17 μM 95% CI). n=3 biological replicates. A non-linear regression was used to calculate the IC_50_ and is shown as the line with the 95% CI included as a watermark. ND = not determined.

To first assess eIF2A function during translation, we titrated recombinant MBP-eIF2A-His6 into mammalian *in vitro* translation reactions using rabbit reticulocyte lysate (RRL) programmed with nanoLuciferase (nLuc) reporter mRNA. RRL is commercially available and is commonly used to test the effects of both wildtype and mutant translation factors when present in excess (36–38), as well providing a robust system to test different *cis-*regulatory elements in mRNAs (32,37,39,40). RRL is also deficient in some translational control factors, providing a “blank slate” to assess how specific factors affect translation. For example, compared to HEK293 cells, RRL naturally contains very low levels of ZNF598, which provided a vital tool to decipher how ZNF598 recognizes collided ribosomes (41). Despite being the source of where eIF2A was initially purified from and identified in, RRL contains low amounts of endogenous eIF2A. Using purified tag-less eIF2A as a standard, we estimate by Western blot that in the subsequent *in vitro* translation reactions that all use 20% RRL (final), the concentration of endogenous eIF2A is ∼2.34 nM (**Supplementary Figure S1B**). By comparison, and by adjusting for the difference in RRL used from published data (36), other initiation factors, specifically eIF4A and eIF4E, are present at ∼740 nM and ∼74 nM, respectively. Thus, we started our titration near the lower end of this range and increased the amount of control protein and eIF2A added. Using purified 80S ribosomes as a standard, we estimate by Western blot for RPS6 (eS6) and RPL7 (uL30) (42) that total ribosome content in 20% RRL is ∼0.95 µM (**Supplementary Figure S1C, D**). Upon separating control *in vitro* translation reactions (kept on ice to prevent translation from proceeding) on traditional sucrose gradients and collecting a continuous A_260_, we used published absorbance conversions (34,35) and determined there is a 3:1:1 ratio of 80S ribosomes (∼0.57 µM), free 40S ribosomal subunits (∼0.19 µM), and free 60S ribosomal subunit (∼0.19 µM), respectively, in 20% RRL. In this RRL *in vitro* translation system, His6-MBP control protein was largely inert in the system (IC_50_ = 14.38 μM with 7.97-40.34 μM 95% CI), but, to our surprise, recombinant eIF2A robustly inhibited translation (IC_50_ = 0.19 μM with 0.14-0.25 µM 95% CI) (**Figure 1B**).

We do not believe the fused MBP tag contributes to inhibition as cleaving off the MBP tag with TEV protease still renders recombinant eIF2A inhibitory (**Supplementary Figure S1E, F**). To ensure this noted inhibition was not due to human eIF2A being expressed in *E. coli* and folding improperly (despite being soluble), we expressed and purified eIF2A from insect Sf9 cells using a baculovirus expression system (**Figure 1C**), which yielded 507 μg from a single 50 mL pellet (an ∼6000-fold increase in yield/mL culture) and repeated the titration in mammalian *in vitro* translation reactions. Insect cell produced recombinant eIF2A robustly inhibited translation (IC_50_ = 0.37 µM with 0.27-0.50 95% CI) (**Figure 1D**). Inhibition did not seem to favor transcripts with a particular sequence or translation efficiency as various reporters containing different 5ʹ UTRs and coding sequences were largely equally repressed upon eIF2A titration (**Supplementary Figure S2**). Human cell (HEK293T) derived recombinant eIF2A-FLAG also repressed translation to a similar degree as insect cell derived recombinant eIF2A (**Supplementary Figure S3A, B**). Additionally, despite control tag proteins being purified in parallel and not being inhibitory, we ruled out the possibility of contaminating RNases influencing the translation reactions by measuring reporter mRNA levels before and after translation. Indeed, reporter mRNA levels were minimally affected after *in vitro* translation in the presence of protein storage buffer, control tag protein, or eIF2A (**Supplementary Figure S3C**).

Even when using TEV protease to cleave affinity tags, a small region of the TEV protease recognition sequence remains, creating a non-native end. As exemplified by recombinant yeast eIF1A and eIF5B (43–46), some recombinant eIFs require native C-termini to function properly. To test whether recombinant tag-less eIF2A with native termini is still inhibitory, we adapted the IMPACT system that uses DTT-inducible inteins and the Sf9 cell expression system to create recombinant human eIF2A with native N- and C-termini (**Figure 1C**). Consistent with tagged-eIF2A (**Figure 1B, D**), tag-less eIF2A robustly inhibited translation (IC_50_ = 0.13 µM with 0.10-0.17 95% CI) (**Figure 1D**).

Several reports have used biochemical and genetic approaches to infer that eIF2A could be able to initiate translation at near-cognate start codons, possibly even at times utilizing Leu-tRNA^Leu^ (15,20,21). Additionally, altered levels of eIFs have been noted to decrease start codon fidelity to favor near-cognate start codons (47,48). Thus, we next asked if eIF2A stimulated or inhibited translation that initiated at near-cognate CUG and GUG start codons. We generated AUG-, CUG-, GUG-, and AAA-encoded 3XFLAG-*Renilla* Luciferase (RLuc) reporters (**Figure 2A** and **Supplementary Figure S4**) that only differed by the start codon and tested how eIF2A affected translation of each. In this set of empirically optimized RLuc reporters (49) (**Supplementary Figure S4**), CUG and GUG start codons were ∼2% as efficient as the canonical AUG start codon *in vitro* (**Figure 2B**), similar to what we have seen for nLuc reporters in perfect Kozak context in HeLa cells (50). As expected, the reporter harboring an AAA start codon, which does not support initiation, was markedly less efficient than both the CUG- and GUG-encoded reporters (**Figure 2B**). Supporting the inhibitory nature of eIF2A we describe above but conflicting the inferred positive role of eIF2A in near-cognate-mediated translation initiation by others, eIF2A produced in either *E. coli* or insect cells equally inhibited translation of AUG-, CUG-, and GUG-encoded RLuc reporter mRNAs (**Figure 2C, D** and **Supplementary Figure S5**). Together, these data reveal a new inhibitory phenotype caused by increased levels of eIF2A.

**Figure 2.**
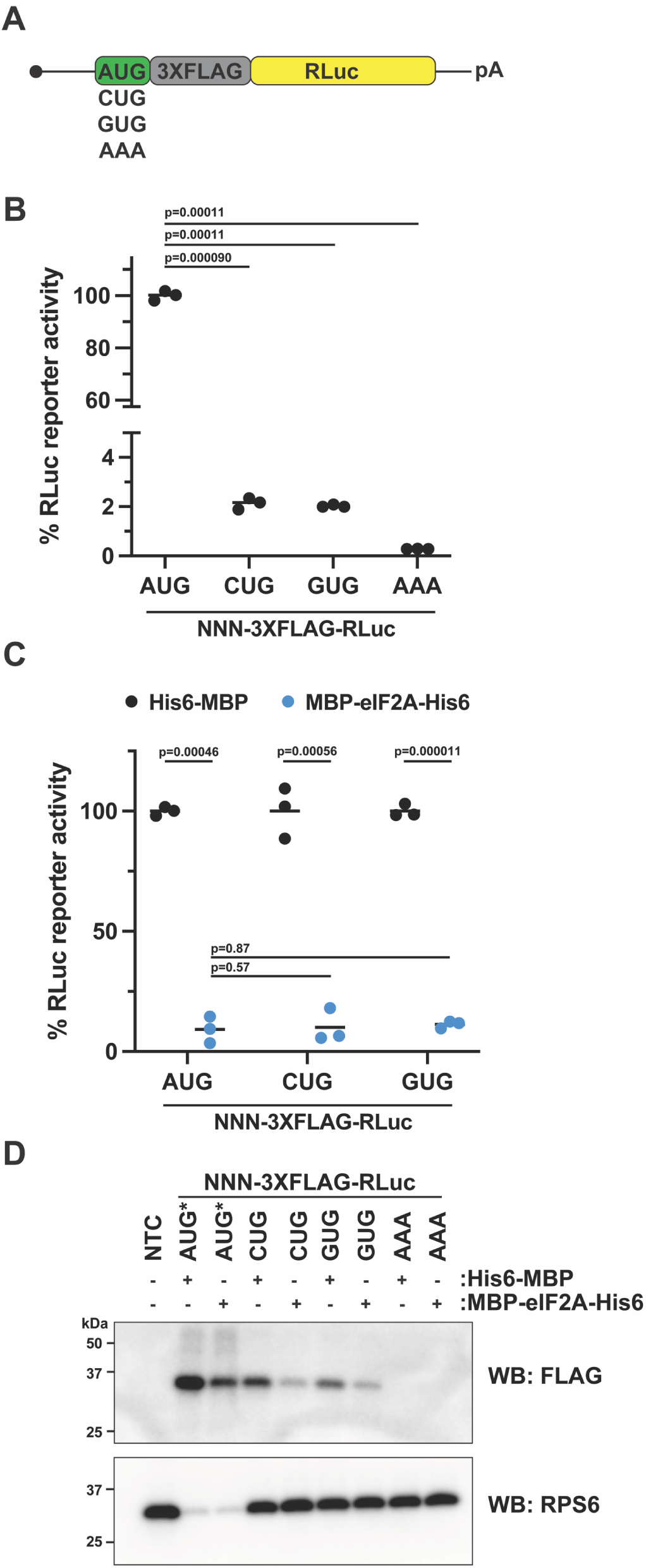
eIF2A inhibits AUG- and near-cognate-initiated translation. A) Diagram of *Renilla* Luciferase (RLuc) reporter mRNAs harboring various start codons and a N-terminal 3XFLAG tag. B) Comparison of AUG- and non-AUG-3XFLAG-RLuc reporter mRNAs translated *in vitro.* Luciferase levels are normalized to AUG-3XFLAG-RLuc. Bars represent the mean. n=3 biological replicates. Comparisons were made using a two-tailed unpaired t-test with Welch’s correction. C) Response of *in vitro* translation reactions programmed with AUG- and non-AUG-3XFLAG-RLuc reporter mRNAs in the presence of 1.68 μM His6-MBP or 1.68 μM MBP-eIF2A-His6. Bars represent the mean. n=3 biological replicates. Comparisons were made using a two-tailed unpaired t-test with Welch’s correction. D) Anti-FLAG Western blot of *in vitro* translation reactions programmed with AUG- and non-AUG-3XFLAG-RLuc reporter mRNAs in the presence of 1.68 μM His6-MBP or 1.68 μM MBP-eIF2A-His6. RPS6 was used as a loading control. AUG* = AUG-3XFLAG-RLuc samples were diluted 1:10 to prevent overexposure of the anti-FLAG blot. NTC = No Template Control.

### Distinct regions of the N- and C-termini are required for eIF2A-mediated translational repression

To better understand how eIF2A inhibits translation, we next sought to determine which elements of eIF2A are required for inhibition. Empirically determined structures for full-length mammalian eIF2A (residues 1-585) are not available, but AlphaFold predicts a globular nine bladed β-propeller at the N-terminus and three alpha helices at the C-terminus connected by flexible linkers (**Figure 3A, left; Supplementary Figure S6A**) (51,52). It should be noted that the predicted flexible linkers are of lower confidence (**Supplementary Figure S6A**). Crystal structures of truncated human eIF2A (residues 4-427; PDB: 8DYS) and truncated *Schizosaccharomyces pombe* eIF2A (residudes 1–424; PDB: 3WJ9) also show a globular nine bladed β-propeller at the N-terminus (53). Kim *et al*. previously reported that eIF2A has three separate functional domains to bind Met-tRNA_i_^Met^, eIF5B, and mRNA (**Figure 3A, right**) (54). Using these defined regions and the AlphaFold predicted structure, we generated a large series of eIF2A deletion mutants (**Figure 3B** and **Supplementary Figure S6B**) and tested how each affected translation of nLuc mRNA (**Figure 3C** and **Supplementary Figure S6C**). Of the 13 mutants, only seven—residues 1-415, 1-430, 1-437, 1-471, 1-480, 416-585, and 533-585— purified to a respectable level of purity with sub-stochiometric levels of GroEL chaperone (**Figure 3B**). However, all seven of these mutants did not inhibit translation (**Figure 3C**). Of the six that did not purify well (**Supplementary Figure S6B**), only 1-503, 1-529, and 1-556 demonstrated some translation inhibition (**Supplementary Figure S6C**). From these data, we designed two new mutants that harbored the complete N-terminus (1–437), a GGS linker, and a portion of the C-terminus (either 504-556 or 529-556) (**Figure 3D**) that purified with low levels of GroEL and inhibited translation at least two-fold (**Figure 3E**). These data support that a single domain in eIF2A is not responsible for translation inhibition.

**Figure 3.**
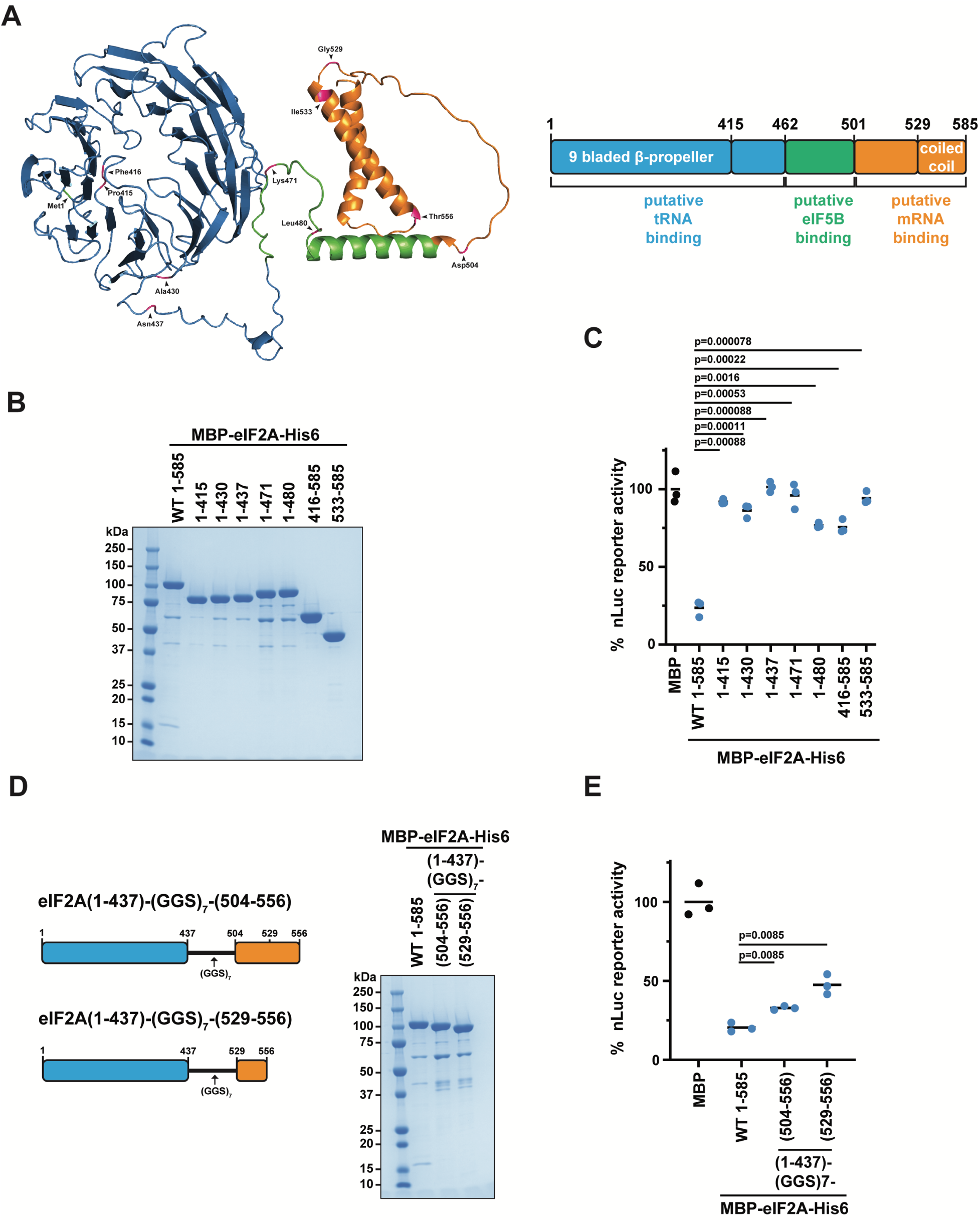
The eIF2A N- and C-termini alone do not inhibit translation. A) AlphaFold structural prediction of full-length human eIF2A with mutation sites from this study colored in pink (left). Schematic of full-length eIF2A with the three previously annotated domains and structural motifs labeled (right). B) SDS-PAGE and Coomassie stain of recombinant WT and mutant MBP-eIF2A-His6. 2 μg of protein was loaded. C) Response of *in vitro* translation reactions programmed with nLuc reporter mRNA in the presence of 1.68 μM His6-MBP, 1.68 μM MBP-eIF2A-His6, or 1.68 μM of the indicated eIF2A mutant. Bars represent the mean. n=3 biological replicates. Comparisons were made using a two-tailed unpaired t-test with Welch’s correction. D) SDS-PAGE and Coomassie stain of the indicated mutant MBP-eIF2A-His6. 2 μg of protein was loaded. E) Response of *in vitro* translation reactions programmed with nLuc mRNA in the presence of 1.68 μM His6-MBP or 1.68 μM MBP-eIF2A-His6 (WT or indicated mutant). Bars represent the mean. n=3 biological replicates. Comparisons were made using a two-tailed unpaired t-test with Welch’s correction.

### eIF2A inhibits translation prior to 48S initiation complex formation independent of initiation factors or initiator tRNA

To decipher which step of translation is being inhibited by eIF2A, we used sucrose gradient ultracentrifugation along with various translation inhibitors to capture and measure the levels of translation complexes at different stages on nLuc reporter mRNA. Lactimidomycin (LTM) is an elongation inhibitor that binds the E-site of 60S subunits and blocks the first 80S translocation step at the start codon (55). Using this inhibitor, we observed that eIF2A decreased 80S formation ∼5-fold on average (**Figure 4A, B** and **Supplementary Figure S7A**). To determine if eIF2A inhibits translation before 80S formation, we repeated the sucrose gradient experiments and trapped 48S initiation complexes (the 43S pre-initiation complex bound to mRNA at start codons) after start codon recognition but before 60S subunit joining by adding the non-hydrolyzable GTP analog, GMPPNP. If eIF2A inhibited translation after start codon recognition, we would expect unchanged levels of 48S complexes. However, our data shows a ∼7-fold decrease in 48S complex levels (**Figure 4C, D** and **Supplementary Figure S7B**). These data support that increased levels of eIF2A inhibit translation prior to start codon recognition.

**Figure 4.**
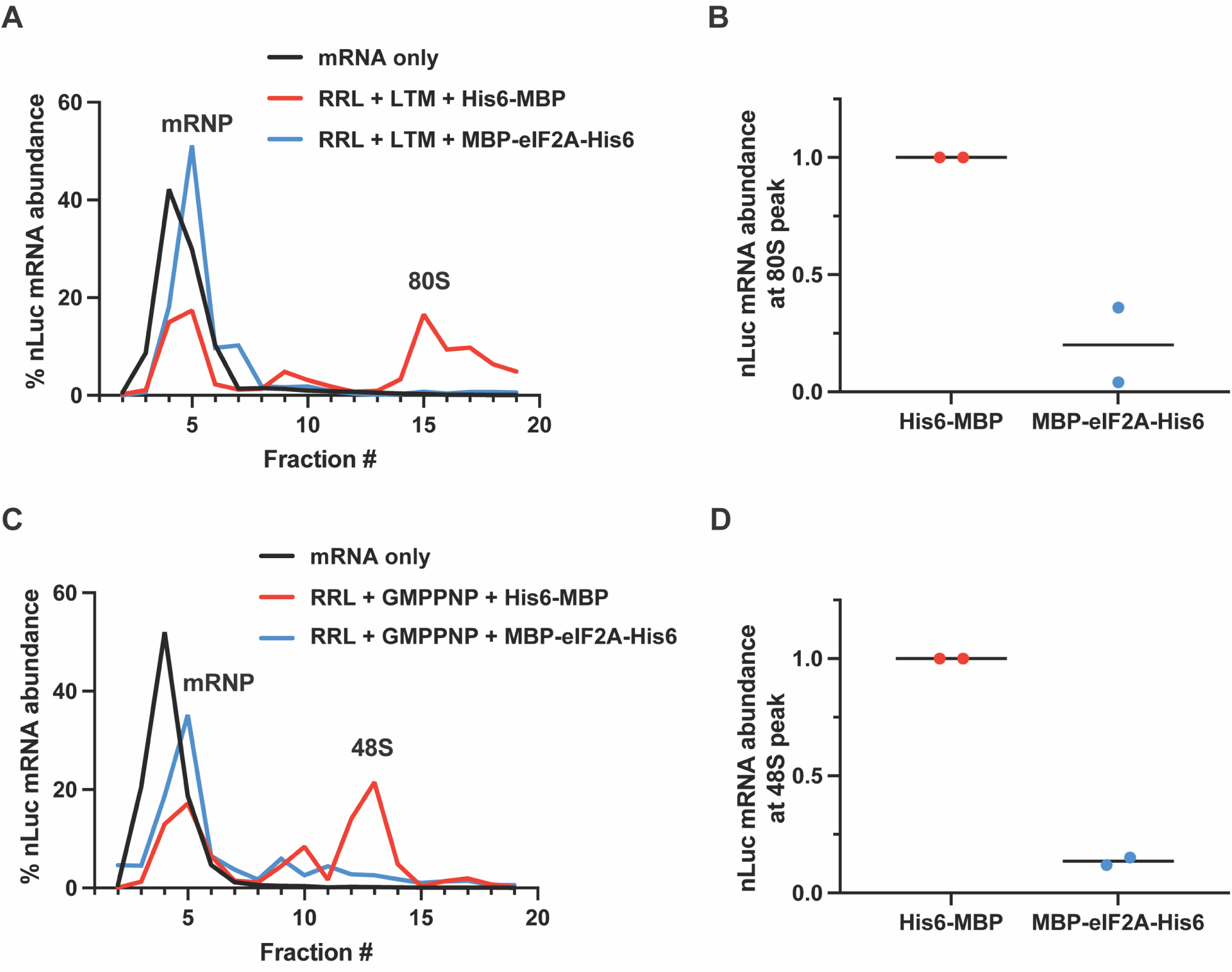
eIF2A inhibits translation prior to 48S initiation complex formation. A) nLuc mRNA distribution along a 5-30% (w/v) buffered sucrose gradient. *In vitro* translation reactions were supplemented with 50 µM lactimidomycin (LTM) to stall 80S ribosomes before the first translocation cycle and with either 1.68 μM His6-MBP or 1.68 μM MBP-eIF2A-His6, then diluted and separated on buffered sucrose gradients. B) Quantification of nLuc mRNA abundance at 80S peak from A. Bars represent the mean. n=2 biological replicates. C) Same as in A, but instead supplemented with 5 mM GMPPNP to capture 48S initiation complexes at the start codon. D) Quantification of nLuc mRNA abundance at 48S initiation complex peak from C. Bars represent the mean. n=2 biological replicates. Replicates are shown in **Supplementary Figure S7**.

To inhibit translation prior to start codon recognition, eIF2A could target initiation factors, the 40S subunit, 43S pre-initiation complex formation and recruitment, or 43S scanning. To decipher among these possibilities, we took advantage of the ability to direct translation using various combinations of initiation factors via viral internal ribosome entry sites (IRESs). Specifically, we used the prototypical type I, II, III, and IV IRESs from poliovirus (PV), encephalomyocarditis virus (EMCV), hepatitis C virus (HCV), and cricket paralysis virus intergenic region (CrPV IGR), respectively (**Figure 5A**) (56–61). Importantly, IRES reporters were A-capped and contained a strong hairpin in the 5’ untranslated region (UTR) to prevent canonical translation and ribosome loading (32). The 3ʹ end of the PV RNA and the PV IRES itself has been shown to be aberrantly translated in RRL without the addition of HeLa cell extract (62,63). Thus, we confirmed that the PV IRES nLuc reporter is translated from the expected start codon of the nLuc reporter ORF by observing a dramatic reduction in reporter signal when the AUG of nLuc was mutated to AAA (**Supplementary Figure S8A, B)**.

**Figure 5.**
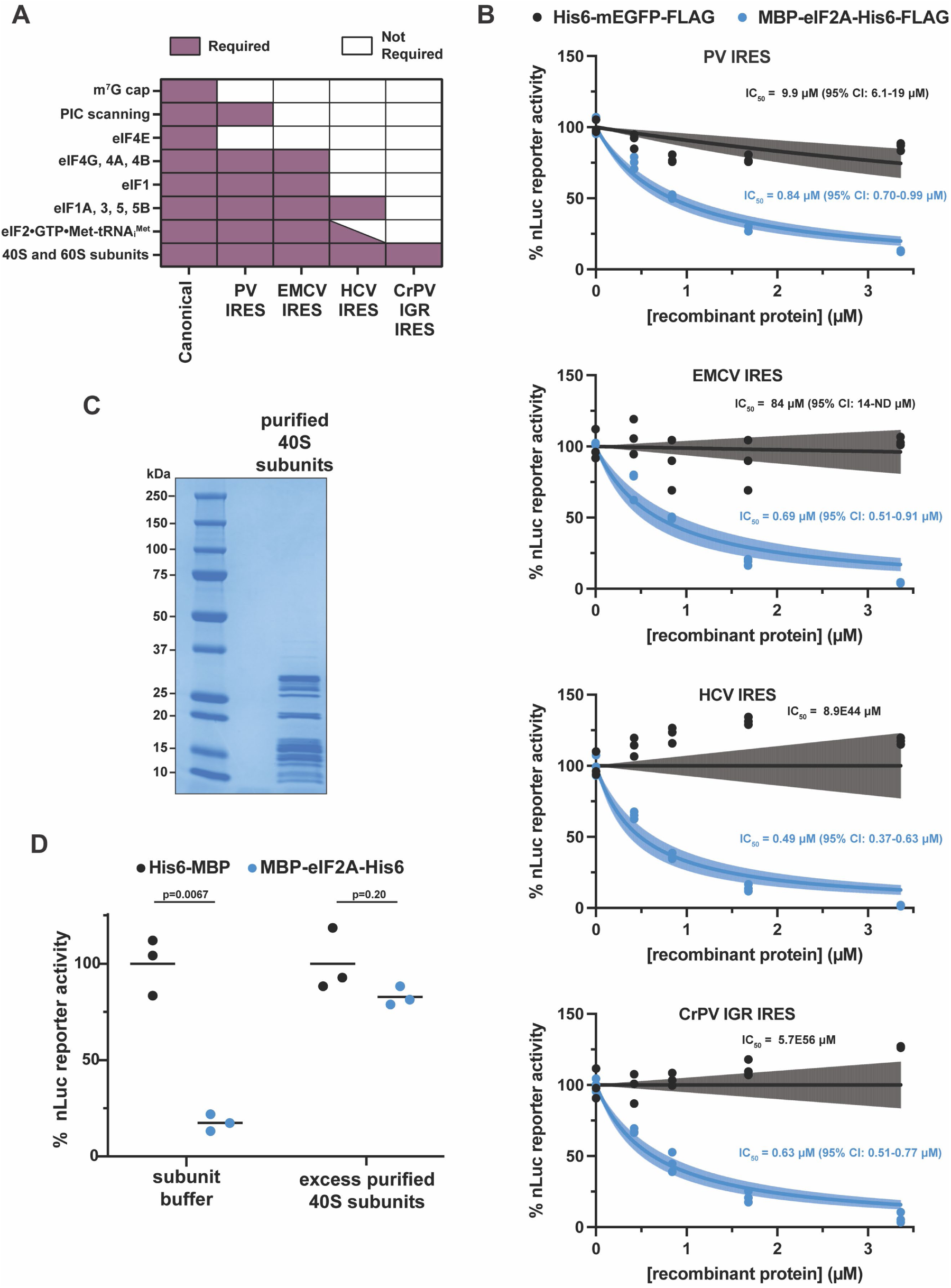
eIF2A inhibits translation independent of eIFs and initiator tRNA. A) Requirements for translation initiation of IRESs types I-IV, which are represented by the poliovirus (PV) IRES, encephalomyocarditis virus (EMCV) IRES, hepatitis C virus (HCV) IRES, and cricket paralysis virus intergenic region (CrPV IGR) IRES, respectively. B) *In vitro* translation of IRES-driven nLuc mRNAs with titration (0-3.4 μM) of either His6-mEGFP-FLAG or MBP-eIF2A-His6-FLAG. IC_50_ values of His6-mEGFP-FLAG were determined for PV IRES (9.9 μM with 6.1-19 μM 95% CI), EMCV IRES (84 μM with 14-ND 95% CI), HCV IRES (8.9E44 μM with +/− infinity μM 95% CI), and CrPV IGR IRES (5.7E66 μM with +/− infinity μM 95% CI) reporters. IC_50_ values of MBP-eIF2A-His6-FLAG were determined for PV IRES (0.84 μM with 0.70-0.99 μM 95% CI), EMCV IRES (0.69 μM with 0.51-0.91 μM 95% CI), HCV IRES (0.49 μΜ with 0.37-0.63 μM 95% CI), and CrPV IGR IRES (0.63 μM with 0.51-0.77 μM 95% CI) nLuc mRNAs. n=3 biological replicates. A non-linear regression was used to calculate the IC_50_ and is shown as the lines with the 95% CI included as a watermark. ND=not determined. C) SDS-PAGE and Coomassie stain of purified 40S ribosomal subunits from RRL. D) Response of *in vitro* translation reactions programmed with nLuc reporter mRNA in the presence of 1.68 μM His6-MBP or 1.68 μM MBP-eIF2A-His6 and supplemented with subunit buffer or 1.35 µM purified 40S ribosomal subunits. Bars represent the mean. n=3 biological replicates. Comparisons were made using a two-tailed unpaired t-test with Welch’s correction.

Upon titration of eIF2A into translation reactions programmed with the four IRES nLuc mRNAs, we observed that recombinant eIF2A inhibited IRES-mediated translation to similar degrees (PV IRES: IC_50_ = 0.84 μM with 95% CI = 0.70-0.99 μM; EMCV IRES: IC_50_ = 0.69 μM with 95% CI = 0.51-0.91 μM; HCV IRES: IC_50_ = 0.49 μM with 95% CI = 0.37-0.63 μM; and CrPV IGR IRES: IC_50_ = 0.63 μM with 95% CI = 0.51-0.77 μM), independent of the subset of eIFs used and did not require 43S scanning (**Figure 5B**). Translation directed from CrPV IGR IRES which supports initiation independent of ribosome scanning, any initiation factors, and the initiator tRNA was also notably inhibited (**Figure 5B**)., suggesting that eIF2A either targets the 40S or 60S subunit.

The CrPV IGR IRES has been shown to bind both the 40S subunit (subsequently recruiting the 60S subunit) and pre-formed vacant 80S ribosomes from salt-washed 40S and 60S subunits (61,64). In both cases, translocation is observed once the 80S is formed on the CrPV IGR IRES. Additionally, RRL is known to contain large amounts of 80S ribosomes over free subunits (**Supplementary Figure S8C**). This raises the possibility that in our assays, the CrPV IGR IRES nLuc mRNA is preferentially interacting with the more abundant 80S ribosomes instead of the 40S subunits first, complicating the readout of the CrPV IGR IRES nLuc reporter. However, we do not believe this possibility is interfering with our interpretations for two reasons. First, cryo-EM studies of native 80S ribosomes from RRL found them to be in states that are translationally inactive or incompatible (e.g., bound by SERBP1•eEF2, bound by IFRD2, or contain tRNA in noncanonical binding sites) (65). Vacant 80S ribosomes in the classic/unrotated state that would mirror the pre-formed ribosomes mentioned above were absent in all samples tested (65). Secondly, in conditions that would allow the IRES to interact with the translation machinery in RRL, we found that the CrPV IGR IRES nLuc mRNA co-sediments with 40S subunits but not 80S ribosomes (**Supplementary Figure S8D**).

Given that 60S subunit joining is downstream from 48S initiation complex formation and eIF2A decreases 48S levels (**Figure 4C, D** and **Supplementary Figure S7B**), we hypothesized that eIF2A is sequestering the 40S subunit, resulting in eIF2A inhibiting translation independent of eIFs or initiator tRNA (**Figure 5A, B**). To test this idea, we supplemented translation reactions with excess purified 40S subunits to rescue translation. In agreement with our hypothesis, excess 40S subunits severely blunted the ability of eIF2A to inhibit translation (**Figure 5C, D**). Together, these data demonstrate that increased levels of eIF2A inhibit translation by sequestering 40S ribosomal subunits.

### eIF2A directly binds the 40S ribosomal subunit

Previous work has shown that eIF2A co-sediments with 40S and 80S ribosomes in yeast and with 40S subunits in HEK293T cells (29,54), but there is limited evidence of eIF2A directly binding to the 40S subunit. Initial work using filter binding assays identified eIF2A could deliver Met-tRNA_i_^Met^ to 40S subunits with an AUG trinucleotide with 5 mM Mg^2+^ (5,10). However, a subsequent report identified eIF2D as a contaminating protein in many eIF2A purifications from RRL and could not reproduce the ability of eIF2A to bind initiator tRNA (6). To determine whether eIF2A interacts with 40S subunits in our translation reactions, we took advantage of the C-terminal His6 tag present on recombinant eIF2A for pulldown assays with Ni^2+^-NTA magnetic beads and then probed by Western blot for RPS6 (eS6) as a marker for 40S subunits. Indeed, RPS6 did co-purify, in a formaldehyde-dependent manner, with MBP-eIF2A-His6 but not with the negative control His6-MBP (**Figure 6A**). We attempted to use deletion mutants harboring only the N-terminal β-propeller (residues 1-415) or only the C-terminal helices (residues 416-585) to test if either terminus was responsible for 40S binding; however, unequal pulldown from the cross-linked samples was observed. Unequal pulldown was also observed with the deletion mutant harboring residues 1-430 (**Supplementary Figure S9A**).

**Figure 6.**
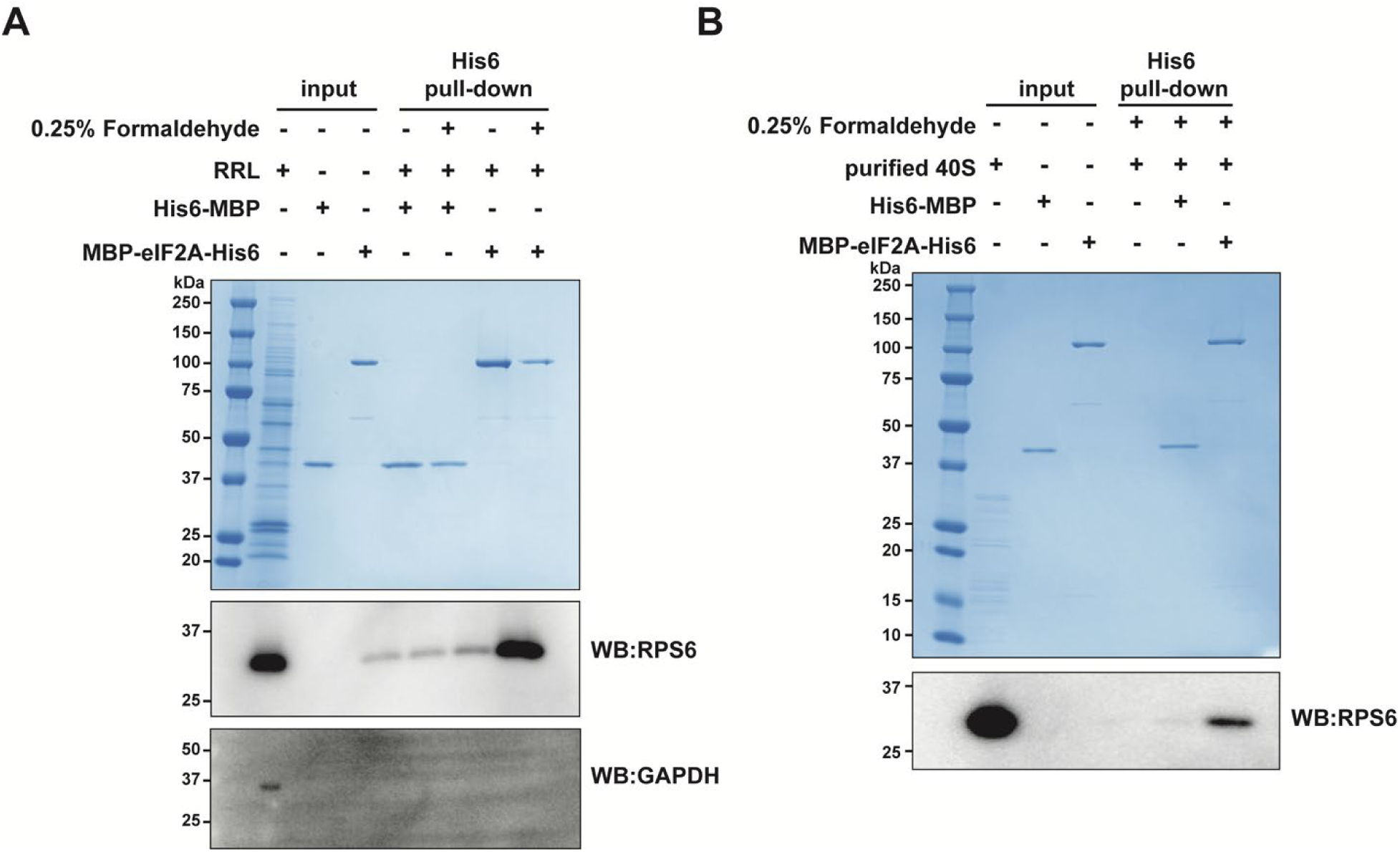
eIF2A directly binds 40S ribosomal subunits. A) SDS-PAGE, Coomassie stain, and Western blot analysis to assess the ability of recombinant His6-MBP or MBP-eIF2A-His6 to pulldown 40S subunits from RRL with Ni^2+^-NTA magnetic beads. 0.25% (v/v) formaldehyde was included in the indicated samples. RPS6 was used as a marker for 40S subunits. GAPDH was used as a negative control. Coomassie staining was used to confirm capture of His6-tagged recombinant proteins. B) Same as in A, but 0.27 μM purified 40S subunits were used instead of RRL. RPS6 was used as a marker for 40S subunits.

The dependence on formaldehyde, a zero-distance crosslinker, suggests a transient but specific association. Although salt levels were at physiological concentrations (i.e., 140 mM KCl) and only non-ionic detergents (i.e., 0.1% (v/v) Triton X-100) were present in the wash buffer, beads were washed rather extensively with five 20 bead volume washes. It should be noted that even high affinity inhibitors that directly bind to ribosomes are completely reversible by wash out. For example, cycloheximide has a reported sub to low µM affinity for the ribosome (55,66) and has an ∼0.1 µM IC_50_ in cells (55), but inhibition is completely reversible upon a change of media in cells (67–69). Nevertheless, to test whether eIF2A directly interacts with 40S subunits, we repeated the His6-pulldowns with purified 40S subunits. Again, RPS6 was co-purified with MBP-eIF2A-His6 but not His6-MBP (**Figure 6B**). Similar results were seen with purified 80S ribosomes, suggesting that eIF2A can interact with the 40S subunit of the 80S ribosome, at least when unbound to mRNA (**Supplementary Figure S9B**). However, we do not believe the ability of eIF2A to bind 80S ribosomes is the cause of inhibition since the decrease in 48S initiation complexes with increased eIF2A levels supports an effect prior to start codon recognition and 80S formation (**Figure 4**). In total, these data support a model in which increased levels of eIF2A inhibit translation initiation by directly binding and sequestering 40S ribosomal subunits (**Figure 7**).

**Figure 7.**
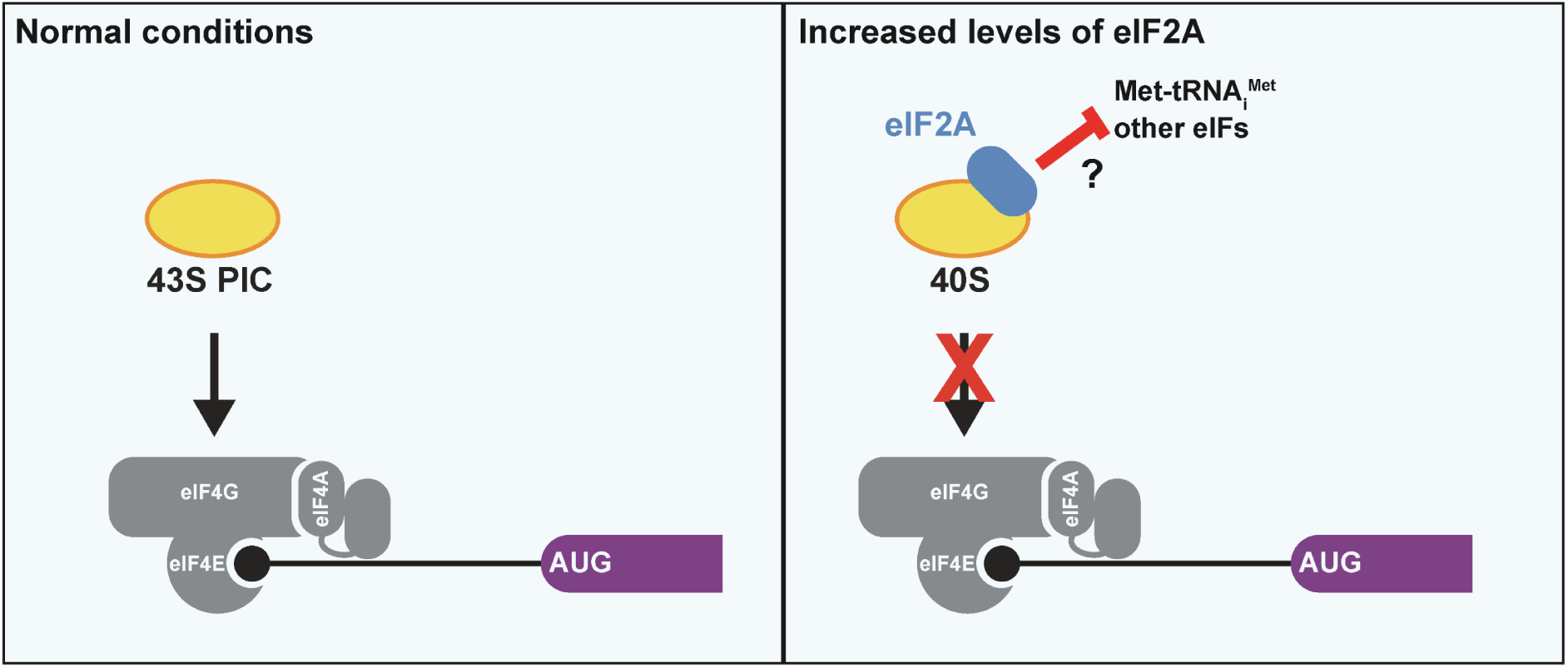
Increased levels of eIF2A inhibit translation by sequestering 40S ribosomal subunits independent of initiation factors and initiator tRNA. Left) In normal conditions, the 43S pre-initiation complex (PIC) is recruited to the 5ʹ end of mRNAs via eIF4F (comprised of eIF4E, eIF4F, and eIF4A). For simplicity, reported eIF2A-dependent translation of specific mRNAs, many of which depend on cell stress, are not shown. Right) Increased levels of eIF2A sequester 40S ribosomal subunits. While eIF2A-mediated inhibition only requires the 40S ribosomal subunit, it remains to be determined if eIF2A blocks or displaces any of the canonical initiation factors or the initiator tRNA.

## DISCUSSION

A correct balance of initiation factors is required for tight control over translation initiation and start codon recognition. For example, overexpression of eIF1 and eIF5 increases and decreases the stringency of start codon selection in cells, respectively (48,70). Here, we show that increased levels of eIF2A inhibit translation initiation independent of eIFs and the initiator tRNA by sequestering 40S ribosomal subunits. This raises the question—how do cells prevent eIF2A from inhibiting translation? eIF2A KO squamous cell carcinoma (SCC) keratinocytes displayed no increases in global translation (measured via ribosome profiling) and no gross change in cell proliferation compared to control SCC keratinocytes (15), suggesting eIF2A does not act as a translation inhibitor during growth in serum-rich media. However, multiple reports demonstrate that eIF2A is non-cytosolic in various cell types, which is highly atypical of translation initiation factors. Thus, eIF2A may naturally be kept in an inactive state by its localization away from the translation machinery. First, Kim *et al*. has reported that in Huh7 cells, eIF2A primarily localizes to the nucleus during typical normal growing conditions but re-localizes to the cytoplasm during cell stress and viral infection (14), possibly to then aid in stress-specific translation. Restraining eIF2A to the nucleus during conditions of high canonical translation could be a robust method for cells to restrict eIF2A from inhibiting translation and sequestering 40S subunits. Whether eIF2A in other cell types share similar localization dynamics has not been reported to our knowledge but would be of value to determine the molecular switch that governs eIF2A re-localization. Nuclear-to-cytoplasmic re-localization during cell stress has been noted for other post-transcriptional regulatory mechanisms and are typically regulated by reversible phosphorylation (71–73). Future work will be required to determine whether eIF2A is reversibly post-translationally modified and regulated in a similar manner during cell stress. Second, Panzhinskiy *et al.* has shown eIF2A to predominantly co-localize with the endoplasmic reticulum (ER) in MIN6 cells (18); however, its localization upon cell stress was not reported when our manuscript was in preparation. It is unclear how eIF2A localized to the ER could play an active role in translation initiation since initiation for all mRNAs occurs in the cytosol. Canonically, translation elongation and termination, not translation initiation, occurs on the ER. Last, and further increasing the complexity of defining a consensus subcellular localization, the Human Protein Atlas shows (with one of two antibodies tested) eIF2A to have a localization pattern consistent of mitochondria in U2OS and A-431 cells (16,17). It remains unreported if primary, non-transformed cells derived from the same tissues as the cancer cell lines listed above demonstrate the same non-cytosolic eIF2A localization. Nevertheless, it will be important for the field to determine if concentrating eIF2A to organelles and away from the translation machinery provides a competitive advantage or increased fitness for cancer cells. Preventing translation inhibition by endogenous eIF2A may be such an advantage.

Although eIF2A most likely does not have a major role in global protein synthesis, as global translation and cell proliferation levels do not appear to be affected upon eIF2A KO in SCC keratinocytes grown in serum-rich media (15) and eIF2A KO mice only develop a phenotype after ∼1 year (30), overexpression and depletion of eIF2A protein levels is reported to affect translation of some specific mRNAs, often during cell stress (14,19,21,22,74). However, such examples are often complicated with either conflicting reports (as described in the Introduction for the HCV IRES), lack of sufficient controls to show specificity (e.g., a negative control reporter that is non-responsive to eIF2A perturbation or confirmation of unchanged mRNA levels), or are not fully supported when the endogenous gene is assayed. For example, protein levels of a reporter mRNA harboring the *BiP* 5ʹ UTR and coding sequence remained steady during ER stress in control cells and decreased ∼30% in eIF2A-knockdown cells, but endogenous BiP protein levels during the same stress conditions increased ∼20% in control cells and remained steady in eIF2A-knockdown cells (19). It will be critical for the field to next provide direct evidence of eIF2A forming initiation complexes on the reported eIF2A-dependent ORFs and mRNAs. Biochemical reconstitution with purified components and selective translation complex profiling (sometimes also referred to as selective 40S profiling) with anti-eIF2A antibodies may provide important insight and more direct evidence of eIF2A forming initiation complexes on specific ORFs and mRNAs.

Enigmatically, the involvement of the Integrated Stress Response (ISR), which revolves around the inhibitory phosphorylation of eIF2α at S51 and the subsequent inhibition of canonical translation initiation, is common among reports that demonstrate eIF2A regulating translation. For example, translation of the subgenomic Sindbis virus 26S mRNA in an eIF2A-dependent manner requires PKR (74), which in turn is activated upon Sindbis virus infection and phosphorylates eIF2α. It is not commonly believed that any of the four mammalian ISR kinases (GCN2, HRI, PERK, PKR) have targets other than eIF2α, at least none that regulate translation initiation since cells that encode the eIF2α-S51A mutation are completely resistant to stress-induced translation inhibition (75,76). A critical objective for the field, and for the documented examples of eIF2A-dependent translation during cell stress, is identifying the driving factor that allows eIF2A to function during cell stress. Is eIF2A activated directly by the ISR kinases or another stress-induced pathway in parallel or downstream? Can eIF2A only successfully compete for the 40S subunit when functional eIF2 levels are low during cell stress? These are not mutually exclusive possibilities, but the latter has been proposed by others (14,74). To our knowledge, only one report has attempted to address these possibilities by testing eIF2A-dependent translation in eIF2α-S51A mutant cells. Tusi *et al*. demonstrated that protein levels of two RAN translation reporters harboring myotonic dystrophy type 2 CCUG•CAGG repeats were markedly reduced in PERK KO cells, PKR KO cells, and eIF2A KO cells, as well as in eIF2α-S51A mutant cells (22). These data would suggest that the observed eIF2A-dependence of the CCUG•CAGG RAN translation reporters is downstream of p-eIF2α. However, in this report, negative control eIF2A-independent reporters to show specificity were lacking and whether the CCUG•CAGG repeats themselves, as it has been shown for other expanded RNA repeats and their translated products (23,77–81), activate the ISR kinases was not addressed. It is imperative for the field to decipher how cell stress, p-eIF2α, and eIF2A are mechanistically connected. We acknowledge that cell stress was not included as a tested variable in the *in vitro* translation reactions with eIF2A in our study; this was due to the inability to efficiently recapitulate cell stress in a cell-free system.

At least two independent reports provide evidence that mammalian cells may contain a buffering system to combat translation inhibition from increased levels of eIF2A. Panzhinskiy *et al*. demonstrated that overexpression of eIF2A in mouse islets via AAV6 delivery and the rat insulin promoter results in a ∼2-fold increase in eIF2A levels and a concatenate decrease of p-eIF2α levels (18). The same gene delivery was able to prevent translation inhibition (measured by polysome abundance) during ER stress treatments; however, whether this effect was due to eIF2A-dependent translation initiation or decreased levels of p-eIF2α from eIF2A overexpression was unaddressed in Panzhinskiy *et al*. when our manuscript was in preparation. A qualitative decrease in p-eIF2α levels is also seen when eIF2A is overexpressed in HEK293T cells (22). These data suggest that cells are capable of sensing eIF2A levels and increase basal levels of initiation by lowering p-eIF2α levels when eIF2A is above normal levels. Whether this buffering effect is from decreased ISR kinase activity, increased PP1•CReP phosphatase activity, and/or increased PP1•GADD34 phosphatase activity is unclear. Mammalian cells have evolved a similar buffering system to recover from cell stress; GADD34 mRNA and protein levels ultimately rise to return p-eIF2α to basal levels. This response is in part due to the hallmark increase in the transcription factor ATF4 that subsequently increases transcription of the *GADD34* gene (82–84). Notably, such buffering systems dependent on gene activation and transcription would be lost in RRL due to reticulocytes naturally being enucleated. Moving forward, the effects of eIF2A levels on p-eIF2α levels (and thus, active eIF2•GTP•Met-tRNA_i_^Met^ ternary complexes) should be examined when investigating eIF2A-dependent translation to rule out possible indirect effects from altering functional ternary complex concentrations.

Previous work reported that residues 462-501 of eIF2A interact with eIF5B, the GTPase that controls subunit joining after start codon recognition, and concluded that eIF2A requires eIF5B to be active in translation during the ISR (54). It is possible that recombinant eIF2A is sequestering eIF5B and subsequently inhibiting a late step of initiation. However, this is likely not the main mechanism of inhibition for many reasons. First, inhibiting eIF5B should have no effect on 48S initiation complex levels when captured by GMPPNP, but eIF2A clearly reduced initiation complex levels (**Figure 4C, D**). Second, eIF2A-mediated inhibition was severely blunted when translation reactions were supplemented with additional 40S subunits, suggesting 40S subunits are being targeted by eIF2A (**Figure 5C, D**). Third, translation initiation directed by the CrPV IGR IRES, which does not require eIF5B, was also inhibited by eIF2A (**Figure 5B**). Lastly, deletion of the residues in eIF2A that were mapped to interact with eIF5B did not prevent inhibition (**Figure 3** and **Supplementary Figure S6B, C**).

Liu *et al.* noted that the nine-bladed β-propeller at the N-terminus of *S. pombe* eIF2A forms a similar structure to that of the central domain of eIF3b (85). In eIF3b, this propeller directly contacts RPS9e (uS4) and eIF3i (85,86). While our experiments did not test if eIF2A displaces eIF3b or other eIF3 subunits from 40S subunits, we do not conclude that this possibility is the primary mode of inhibition due to the fact that the CrPV IGR IRES was susceptible to eIF2A inhibition (**Figure 5B**). Additionally, multiple deletion eIF2A mutants with the N-terminal domain alone (residues 1-415, 1-430, and 1-437) did not inhibit translation (**Figure 3C**). Instead, our mutational analysis testing different eIF2A domains reveals that a single domain on its own is not able to inhibit translation (**Figure 3**). Initial work described eIF2A as being able to deliver initiator tRNA to the 40S subunit only when an AUG trinucleotide was present (5,10,11), raising the possibility that eIF2A delivered initiator tRNA to the P site during start codon recognition. However, the exact positioning of eIF2A on the 40S subunit with or without initiator tRNA is still unknown. Future structural work investigating an eIF2A•40S ribosomal subunit complex will be critical to further define how eIF2A contributes to start codon recognition. A major hurdle of recovering a high yield of recombinant eIF2A has now been lifted as we demonstrate here both bacterial- and insect cell-synthesized human eIF2A are active and recombinant eIF2A interacts with 40S ribosomal subunits independent of any additional factors.

## DATA AVAILABILITY

The data underlying this article are available in the article and in its online supplementary data.

## ACKNOWLEDGEMENTS

Experiments were conceived and performed by DJG and MGK with input and help from DJL. The manuscript was written by DJG and MGK with input from DJL. We thank members of the Kearse lab for critically reading the manuscript and Kurt Fredrick, Guramrit Singh, and Eugene Valkov for stimulating conversations. We also thank The Ohio State Comprehensive Cancer Center Genomics Shared Resource (OSUCCC GSR), which is supported by NIH grant P30CA016058, for help with Sanger sequencing.

---

**SUPPLEMENTARY DATA – Grove et al.**

**Supplementary Table S1.** Values for heat map in Supplementary Figure S1.

| Cell Line | Tissue/Cell Type | eIF2α (ppm) | eIF2β (ppm) | eIF2γ (ppm) | eIF2A (ppm) |
| --- | --- | --- | --- | --- | --- |
| HeLa | Cervical | 213 | 204 | 110 | 198 |
| A549 | Lung | 214 | 196 | 109 | 106 |
| HEK293 | Kidney | 182 | 178 | 92.5 | 167 |
| MCF7 | Breast | 206 | 191 | 102 | 190 |
| U2OS | Bone | 204 | 181 | 96.7 | 229 |
| Jurkat | T lymphocyte | 217 | 213 | 109 | 250 |
| LnCap | Prostate | 202 | 194 | 102 | 185 |
| RKO | Colon | 233 | 208 | 117 | 210 |
| HepG2 | Liver | 226 | 213 | 114 | 202 |

**Supplementary Table S2.**
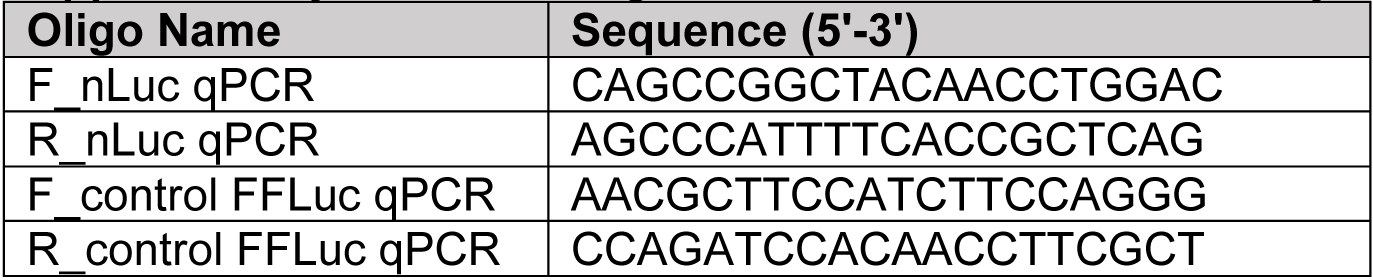
Oligonucleotides used in this study.

**Supplementary Figure S1.**
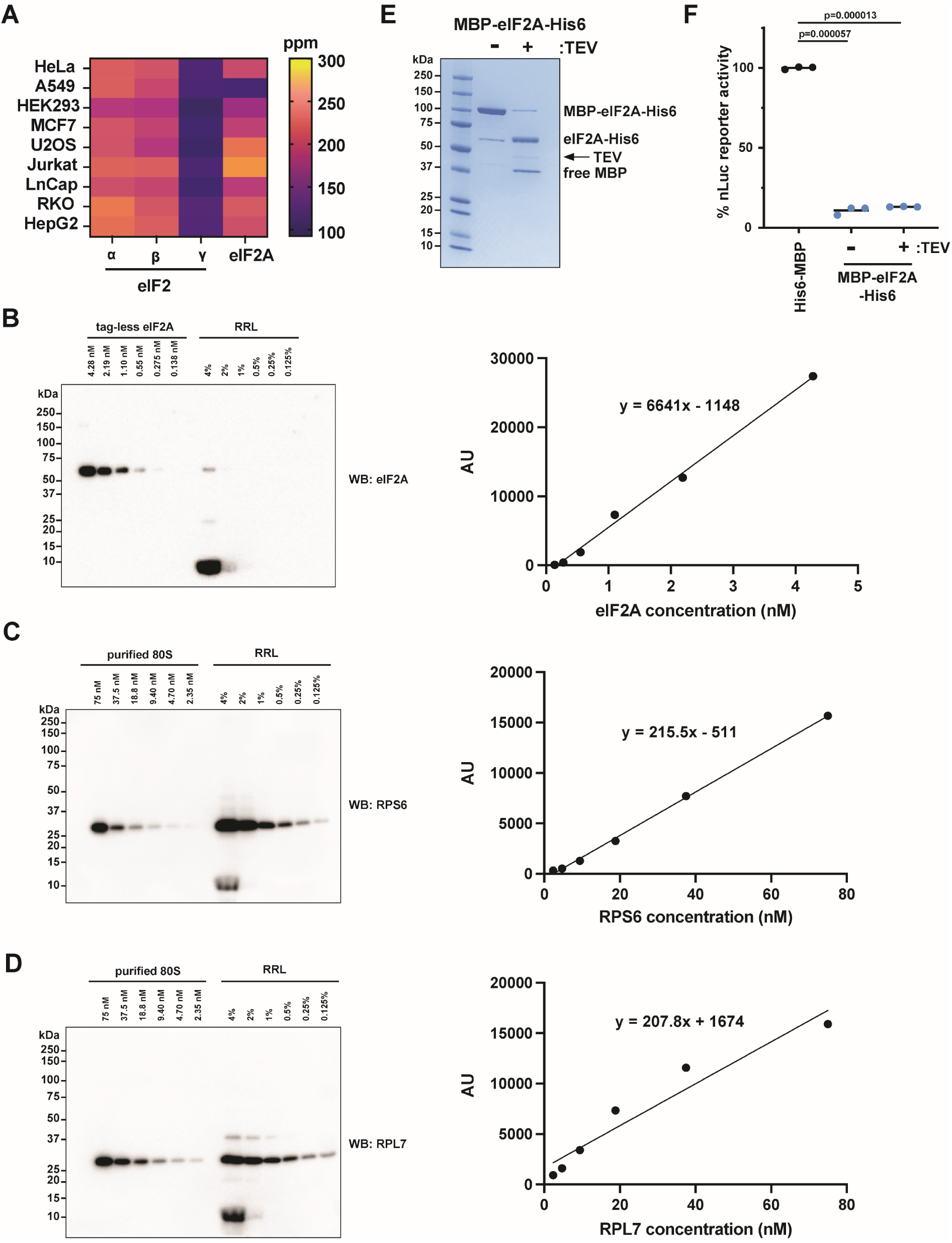
eIF2(α,β,γ) and eIF2A are expressed at similar levels in various human cell types and eIF2A-mediated inhibition remains with TEV protease cleavage. A) Comparison of endogenous eIF2(α, β, γ) and eIF2A protein levels in various human cell types. Data was obtained from pax-db.org using the Geiger, MCP, 2012 dataset. Ppm = parts per million. It should be noted that pax-db.org also contains data sets calculated from spectral counts (these data sets have “SC” in their name) that are less confident due to the lower protein coverage. B-D) Calculation of eIF2A (B), RPS6 (C), and RPL7 (D) concentrations in RRL by Western blot. Recombinant tag-less eIF2A and purified 80S ribosomes from RRL were titrated as a standard curve; quantification showed signal was in the linear dynamic range. The intensity of the 4% RRL sample was used for eIF2Α and the 1% RRL sample was used for RPS6 and RPL7 in the line equation and then multiplied by a dilution factor of 5 or 20, respectively, to determine the concentration of each protein in 20% RRL. For RPS6 and RPL7, these concentrations were averaged since 40S and 60S ribosomal subunits are equimolar. E) SDS-PAGE and Coomassie stain of MBP-eIF2A-His6 without and with TEV protease treatment. F) Response of *in vitro* translation reactions programmed with nLuc mRNA supplemented with 1.68 μM mock-cleaved and 1.68 μM TEV protease-cleaved MBP-eIF2A-His6. Bars represent the mean. n=3 biological replicates. Comparisons were made using a two-tailed unpaired t-test with Welch’s correction.

**Supplementary Figure S2.**
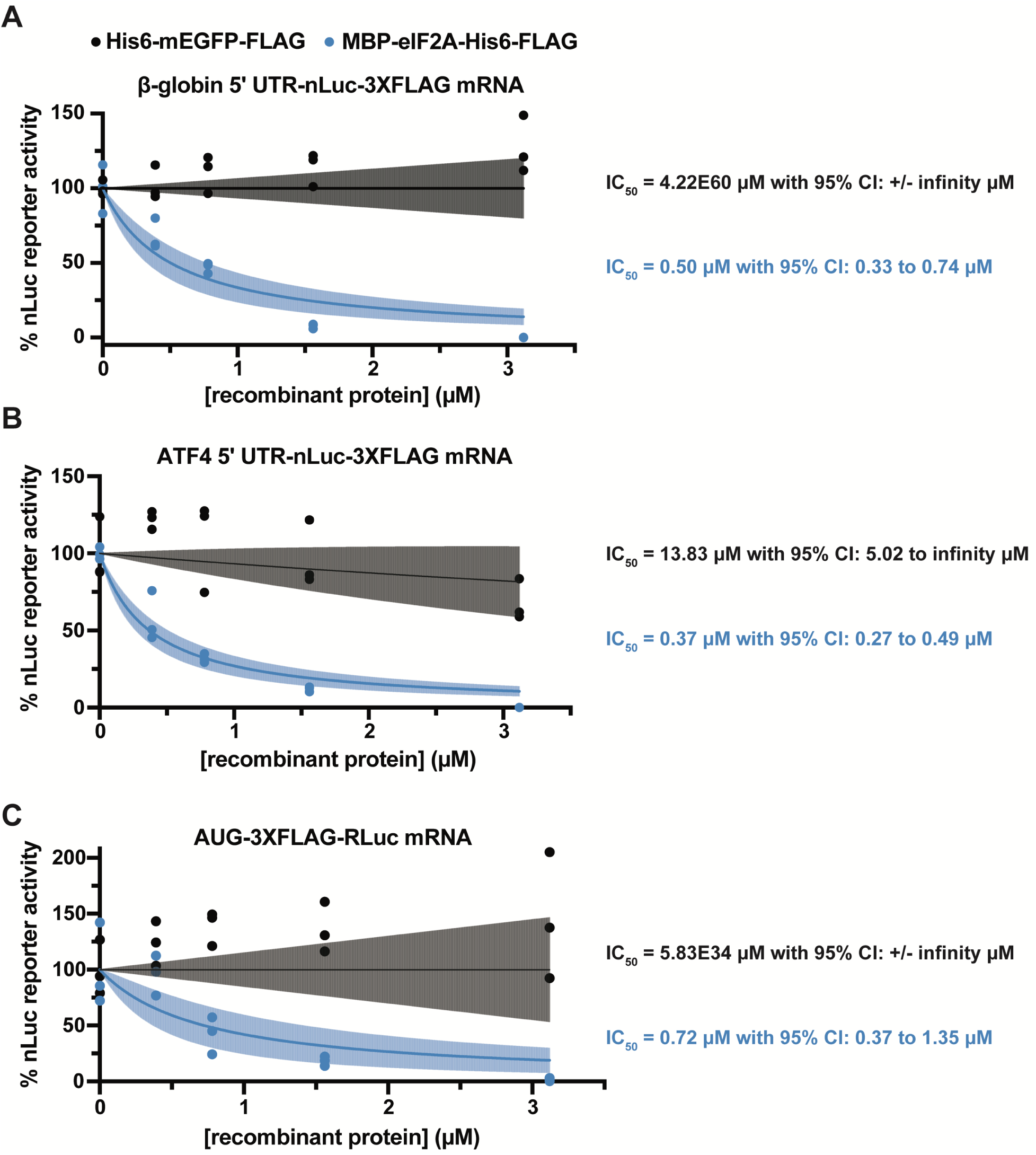
Recombinant eIF2A inhibits translation of mRNAs with different 5ʹ UTRs and coding sequences. A-C) *In vitro* translation of different reporter mRNAs with a titration (0-3.12 μM) of His6-mEGFP-FLAG or MBP-eIF2A-His6-FLAG. mRNAs tested were β-globin 5ʹ UTR nLuc mRNA (A), ATF4 5ʹ UTR nLuc mRNA (B), and AUG-3XFLAG-RLuc mRNA (C). n=3 biological replicates. A non-linear regression was used to calculate the IC_50_ and is shown as the line with the 95% CI included as a watermark.

**Supplementary Figure S3.**
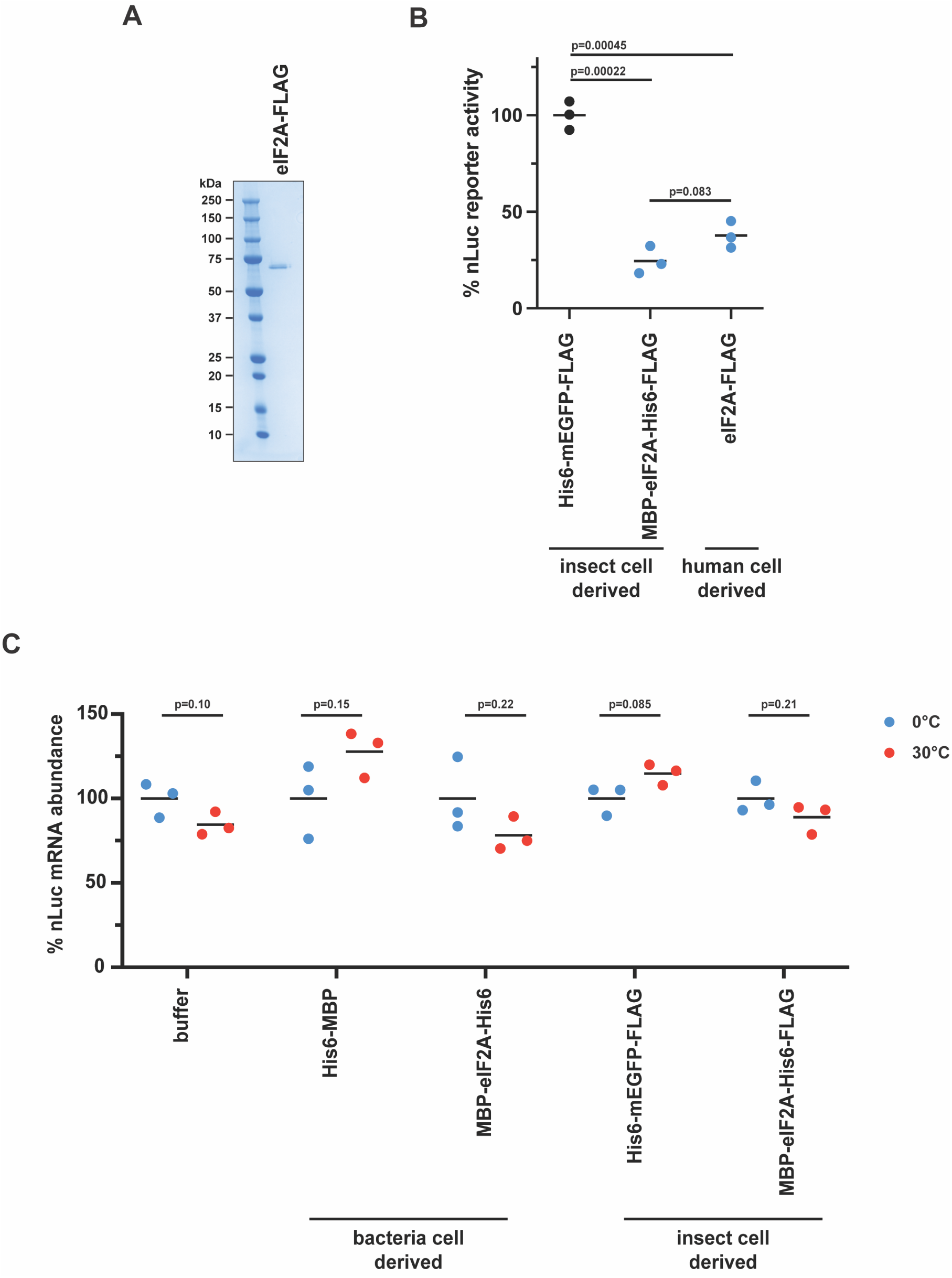
Human cell derived eIF2A-FLAG is inhibitory and recombinant eIF2A does not alter reporter mRNA levels during translation. A) SDS-PAGE and Coomassie stain of recombinant eIF2A-FLAG expressed and purified from HEK293T cells (obtained from OriGene). 1 µg was loaded. B) Response of *in vitro* translation reactions programmed with nLuc reporter mRNAs in the presence of 1.2 μM insect cell derived His6-mEGFP-FLAG, 1.2 μM insect cell derived MBP-eIF2A-His6-FLAG, or 1.2 μM human cell derived eIF2A-FLAG. Bars represent the mean. n=3 biological replicates. Comparisons were made using a two-tailed unpaired t-test with Welch’s correction. C) Relative levels of nLuc reporter mRNA before and after translation with Protein Storage Buffer, 1.68 μM *E. coli* derived His6-MBP, 1.68 μM *E. coli* derived MBP-eIF2A-His6, 1.68 μM insect cell derived His6-mEGFP-FLAG, or 1.68 μM insect cell derived MBP-eIF2A-His6-FLAG. Separate identical reactions were either left on ice (0°C) or translated at 30°C for 30 min. nLuc mRNA abundance for each condition was relative to 0°C samples kept on ice. Reactions were spiked with 0.2 ng control FFLuc mRNA before total RNA was extracted using TRIzol. cDNA was subsequently synthesized and analyzed by RT-qPCR. Bars represent the mean. n=3 biological replicates. Comparisons were made using a two-tailed unpaired t-test with Welch’s correction.

**Supplementary Figure S4.**
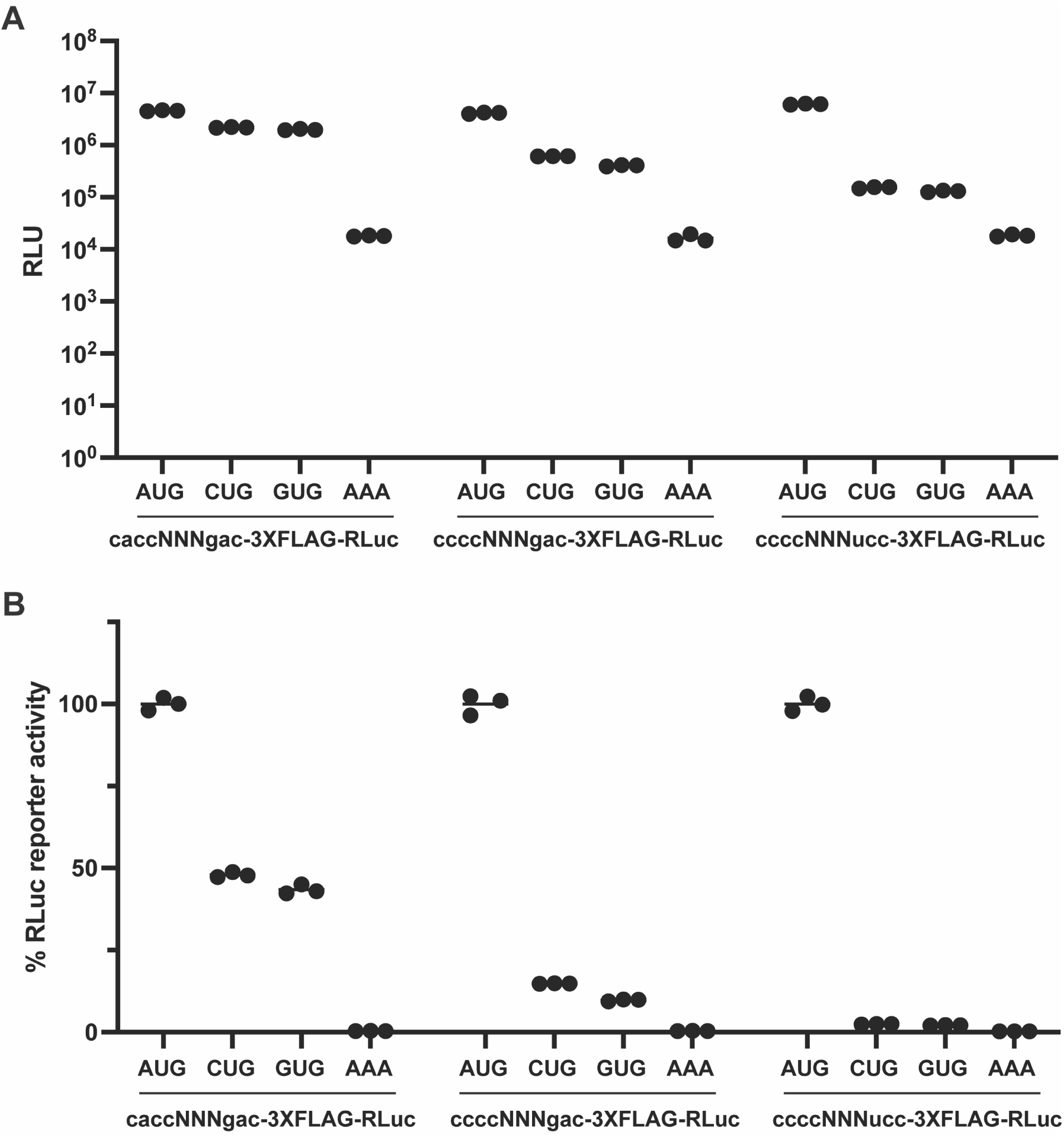
Optimization of RLuc reporter mRNAs. A) *In vitro* translation of nLuc reporter mRNA set 1 (caccNNNguc-3XFLAG-RLuc), set 2 (ccccNNNguc-3XFLAG-RLuc), or set 3 (ccccNNNucc-3XFLAG-RLuc) in RRL. The Kozak sequence surrounding the AUG or non-AUG start codon is perfect (set 1) or mutated (sets 2 and 3). Wei *et al*. has shown that imperfect Kozak sequences cause more drastic efficiencies between AUG and near-cognate start codons in RRL (49). Raw luciferase values (relative luciferase units; RLU) are reported. Bars represent the mean. n=3 biological replicates. B) Same as in A, but each reporter set is relative to the respective AUG-encoded reporter. Bars represent the mean. n=3 biological replicates. Set 3 reporters are used in Figure 2 and **Supplementary Figure S5**. The AUG-RLuc in set 1 is used in **Supplementary Figure S2C**. Complete sequences of these reporters are provided in the **Supplementary Data**.

**Supplementary Figure S5.**
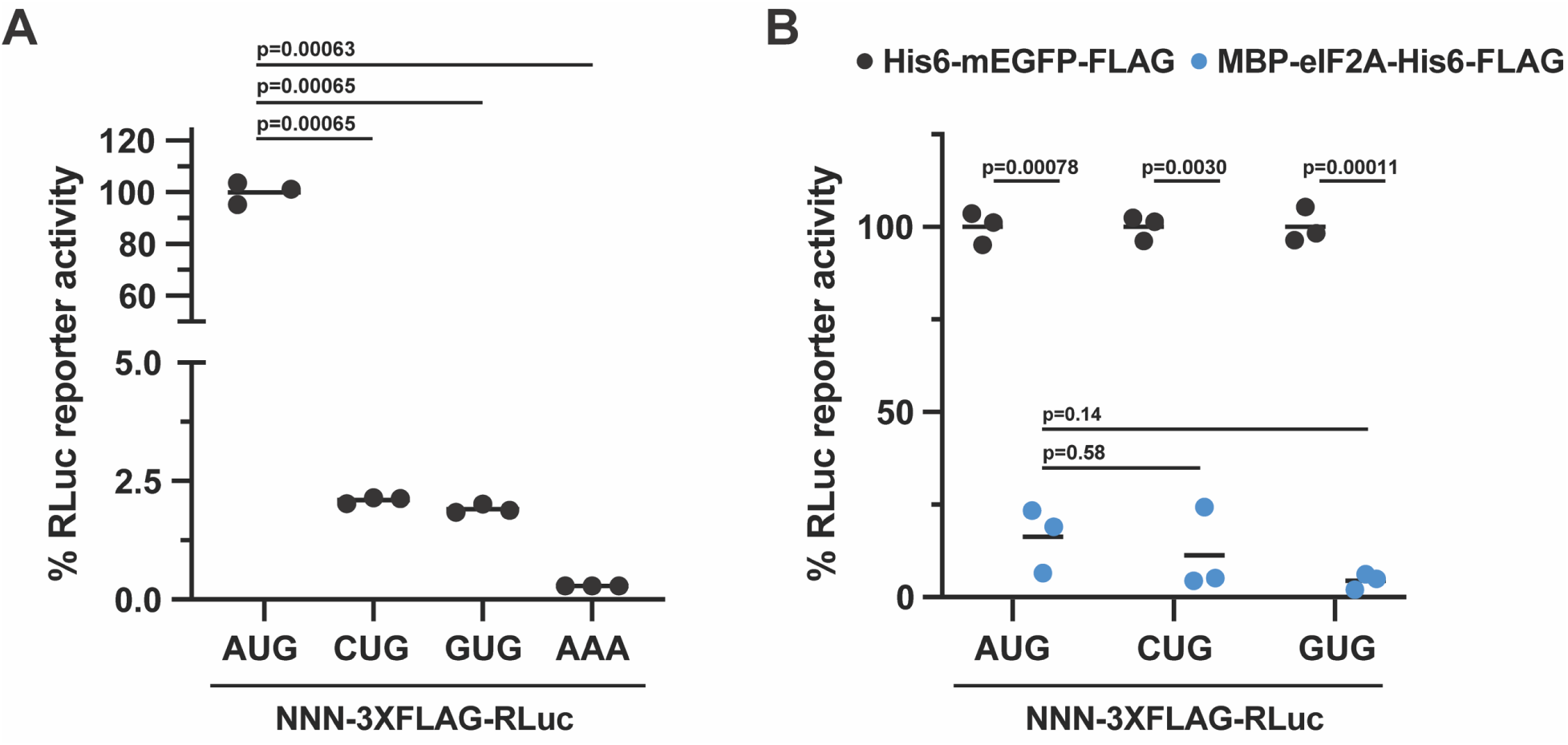
Insect cell derived eIF2A inhibits AUG- and non-AUG translation. A) Comparison of AUG- and non-AUG-3XFLAG-RLuc reporter mRNAs translated *in vitro* in the presence of 1.68 μΜ His6-mEGFP-FLAG. Set 3 RLuc reporters from **Supplementary Figure S4** were used. Luciferase levels are normalized to AUG-3XFLAG-RLuc. Bars represent the mean. n=3 biological replicates. Comparisons were made using a two-tailed unpaired t-test with Welch’s correction. B) Response of *in vitro* translation reactions programmed with AUG- and non-AUG-3XFLAG-RLuc reporter mRNAs in the presence of 1.68 μM His6-mEGFP-FLAG or 1.68 μM MBP-eIF2A-His6-FLAG. Bars represent the mean. n=3 biological replicates. Comparisons were made using a two-tailed unpaired t-test with Welch’s correction.

**Supplementary Figure S6.**
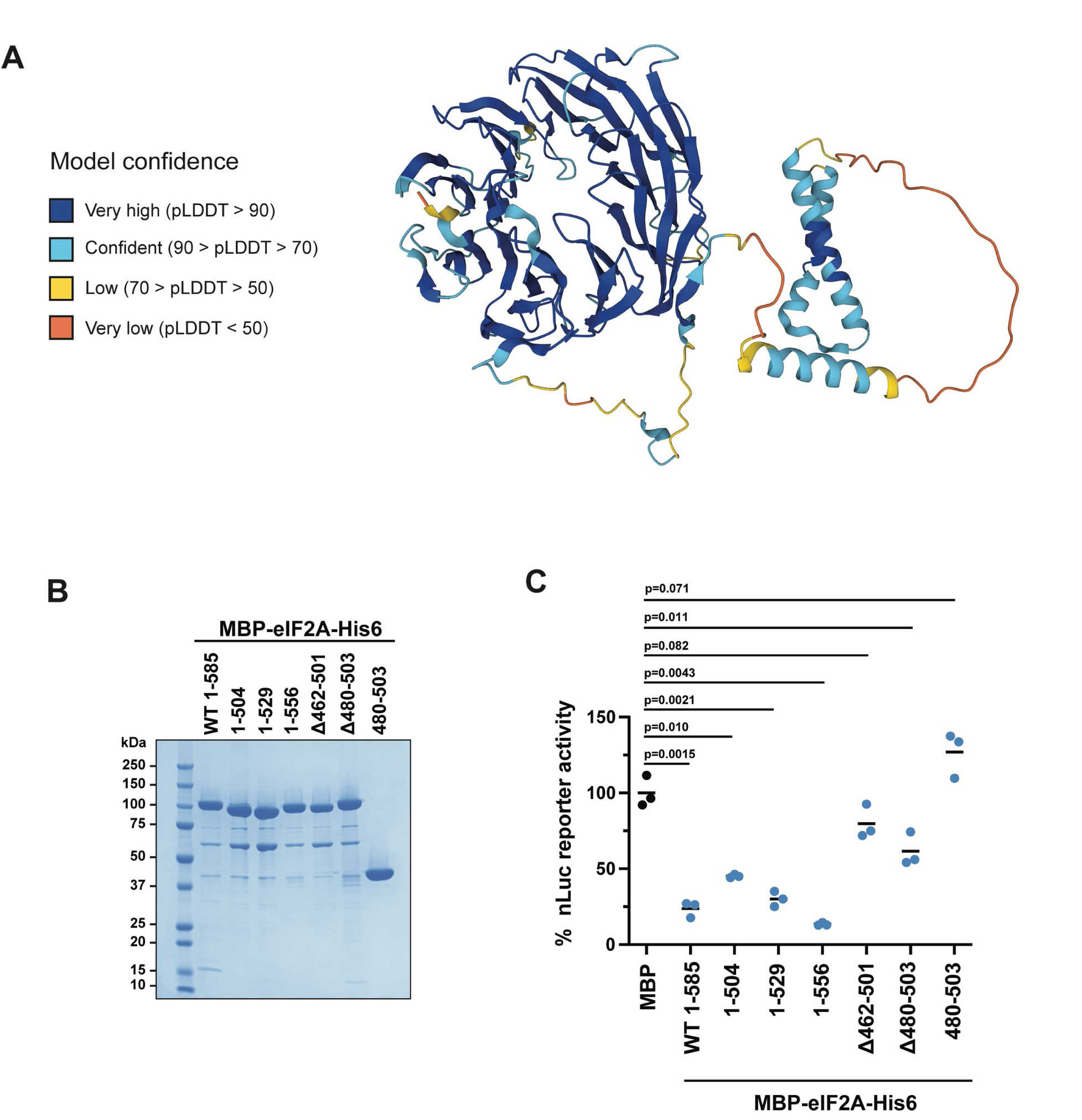
Single domain mutants of eIF2A are not translationally repressive. A) AlphaFold structural prediction of full-length human eIF2A with confidence coloring, which uses the predicted local distance difference test (pLDDT). B) SDS-PAGE and Coomassie stain of recombinant MBP-eIF2A-His6 mutants. 2 μg of protein was loaded. C) Response of *in vitro* translation reactions programmed with nLuc mRNA in the presence of 1.68 μM of the indicated recombinant protein. Bars represent the mean. n=3 biological replicates. Comparisons were made using a two-tailed unpaired t-test with Welch’s correction.

**Supplementary Figure S7.**
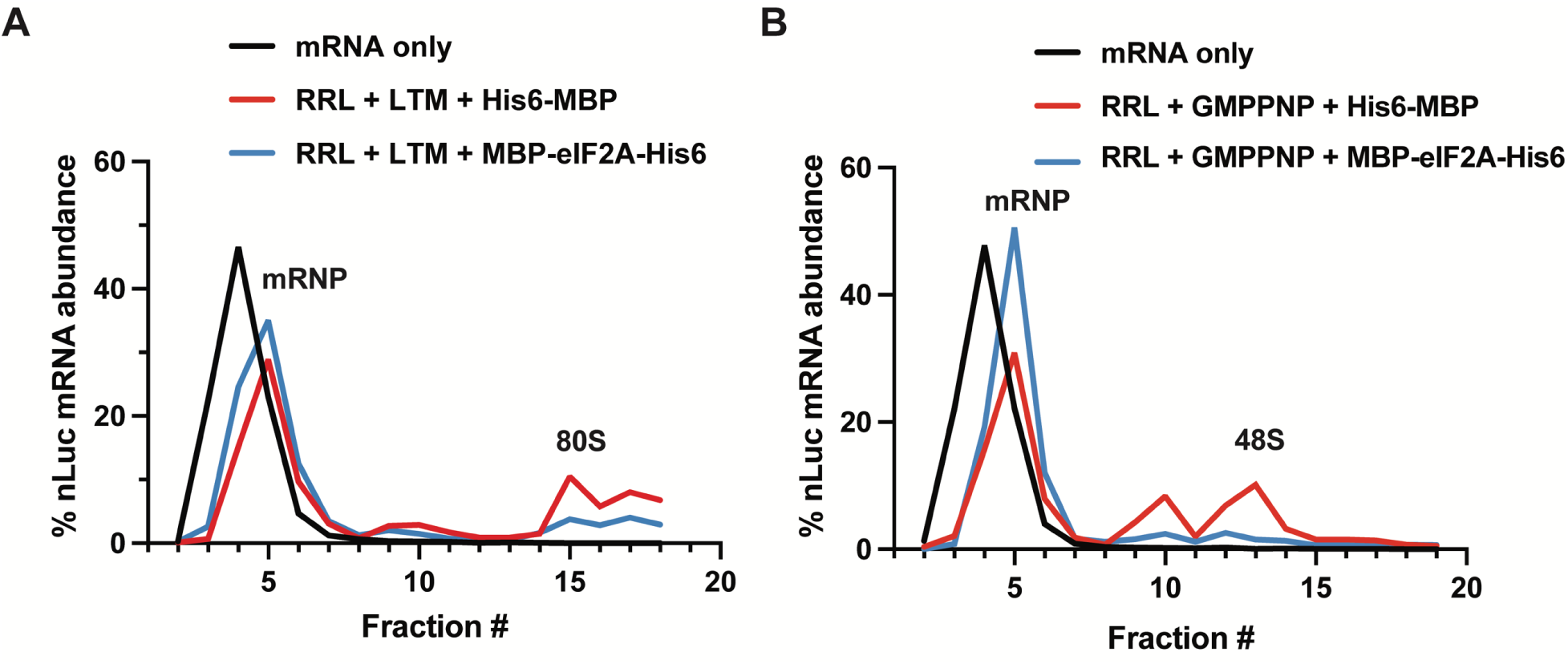
eIF2A represses translation initiation before 48S initiation complex formation. Replicate gradients as shown in Figure 4. A) nLuc mRNA distribution along a 5-30% (w/v) buffered sucrose gradient. *In vitro* translation reactions were supplemented with 50 µM lactimidomycin (LTM) to stall 80S ribosomes before the first translocation cycle and with either 1.68 μM His6-MBP or 1.68 μM MBP-eIF2A-His6, then diluted and separated on buffered sucrose gradients. B) Same as in A, but instead supplemented with 5 mM GMPPNP to capture 48S initiation complexes at the start codon.

**Supplementary Figure S8.**
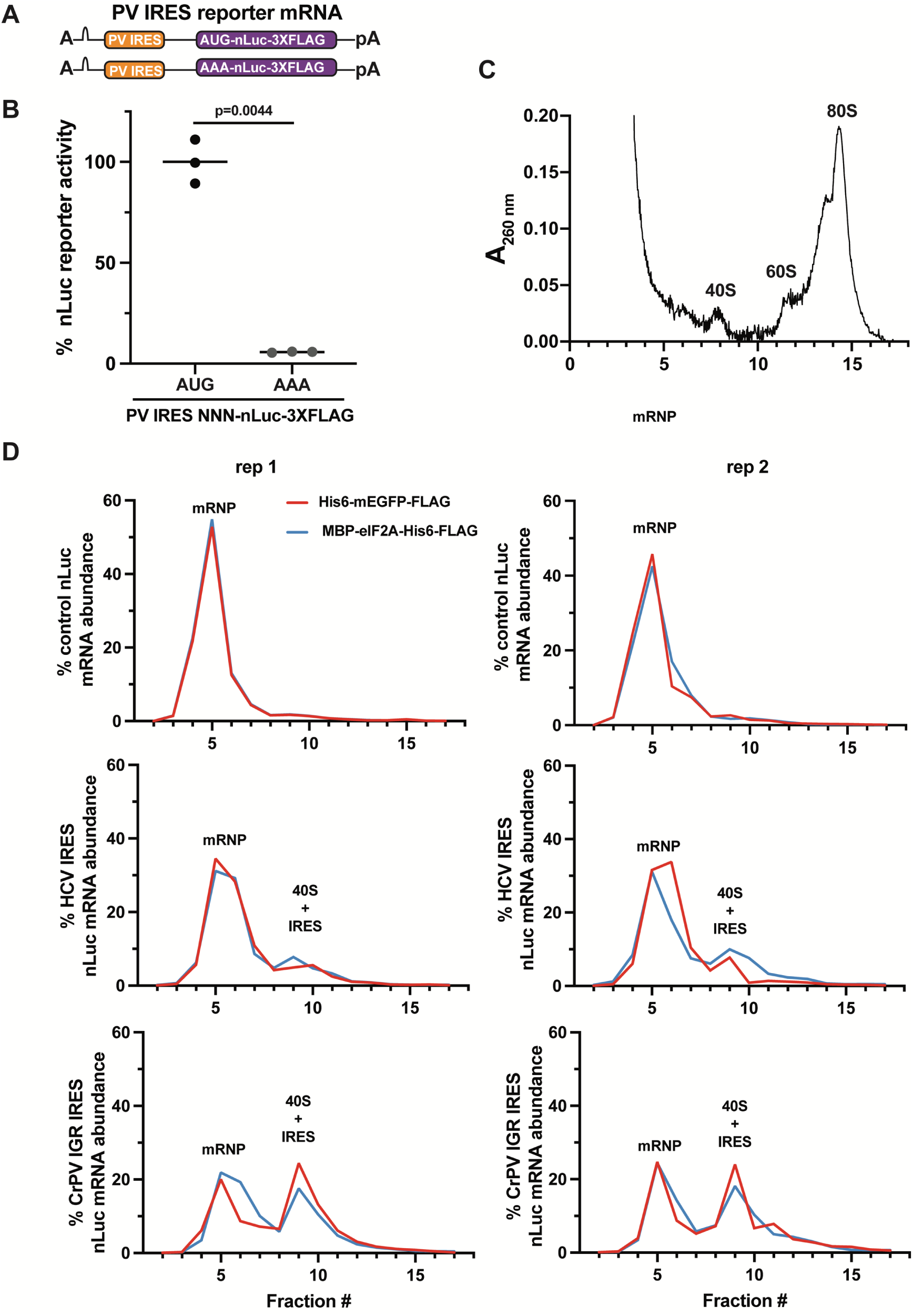
Control experiments with PV, HCV, and CrPV IGR IRES nLuc mRNA reporters. A) Schematic of PV IRES nLuc reporters with the nLuc ORF harboring an AUG or AAA start codon. B) Comparison of *in vitro* translation reactions programmed with either PV IRES nLuc mRNA or PV IRES (AUG to AAA) nLuc mRNA. The AUG to AAA mutation dramatically reduced the signal produced from the reporter, providing evidence that the desired AUG start codon of nLuc is primarily being used in RRL. Bars represent the mean. n=3 biological replicates. Comparisons were made using a two-tailed unpaired t-test with Welch’s correction. C-D) The ability of nLuc mRNA, HCV IRES nLuc mRNA, and CrPV IGR IRES nLuc mRNA to interact with 40S ribosomal subunits and 80S ribosomes in RRL was assessed by 5-30% (w/v) sucrose gradient ultracentrifugation. Translation reactions were assembled identically as in Figure 5B with control (no IRES) nLuc mRNA, HCV IRES nLuc mRNA, or CrPV IGR IRES nLuc mRNA in the presence of 1.68 μM His6-mEGFP-FLAG or 1.68 μM MBP-eIF2A-His6-FLAG but incubated on ice for 30 min to allow interactions between the mRNAs and translation machinery (samples were *NOT* incubated at 30°C as in Figure 5B). Reactions were then diluted and separated on buffered sucrose gradients, and reporter mRNA abundance was measured across the gradient as described in the Materials and Methods. Duplicates are shown side-by-side. The A_260 nm_ trace of an untranslated RRL reaction (assembled and kept on ice, *NOT* incubated at 30°C) is shown in C. Control (no IRES) nLuc mRNA did not co-sediment with 40S subunits. A significant proportion of the HCV IRES nLuc mRNA and CrPV IGR IRES nLuc mRNA did sediment in 40S-containing fractions, with a higher proportion of CrPV IGR IRES nLuc mRNA being found in the 40S-containing fractions than HCV IRES nLuc mRNA. These data demonstrate that these assay conditions are favorable for both IRESs to stably interact with the 40S subunit. Despite the CrPV IGR IRES nLuc mRNA being able to bind pre-formed vacant 80S ribosomes from salt-washed 40S and 60S subunits (64), little to no CrPV IGR IRES nLuc mRNA sedimented in the 80S-containing fractions. Interestingly, although inhibiting translation of the HCV IRES and CrPV IGR IRES nLuc reporter mRNAs (Figure 5B), addition of recombinant eIF2A did not robustly prevent both IRES nLuc mRNAs from co-sedimenting with 40S subunits. Nor did eIF2A cause more IRES nLuc mRNAs to accumulate at the top of the gradient as would be expected if eIF2A was interfering with 40S subunits interacting with either IRES. To speculate, these data overall may suggest that eIF2A is preventing mRNA from being stably inserted into the mRNA channel in the 40S subunit. Unpublished eCLIP data for eIF2A from Wei *et al*. provides some evidence that eIF2A crosslinks to rRNA in the vicinity of the mRNA exit channel (87). The positioning/structure of eIF2A on the 40S is not empirically determined in Wei *et al.* when our manuscript was in preparation or in any other manuscript that we know of. Even for IRESs, which bind to the E, P, and/or A sites of the ribosome first, the mRNA must still be inserted into the mRNA channel after initiation to efficiently allow the ribosome to decode and translocate along the coding sequence. In canonical cap- and scanning-dependent translation, eIF4F bound to the mRNA recruits the 43S PIC and the mRNA is then loaded into the mRNA channel of the 40S subunit; however, this complex is not stable like IRES•40S complexes until start codon recognition occurs and the canonical 48S initiation complex is formed.

**Supplementary Figure S9.**
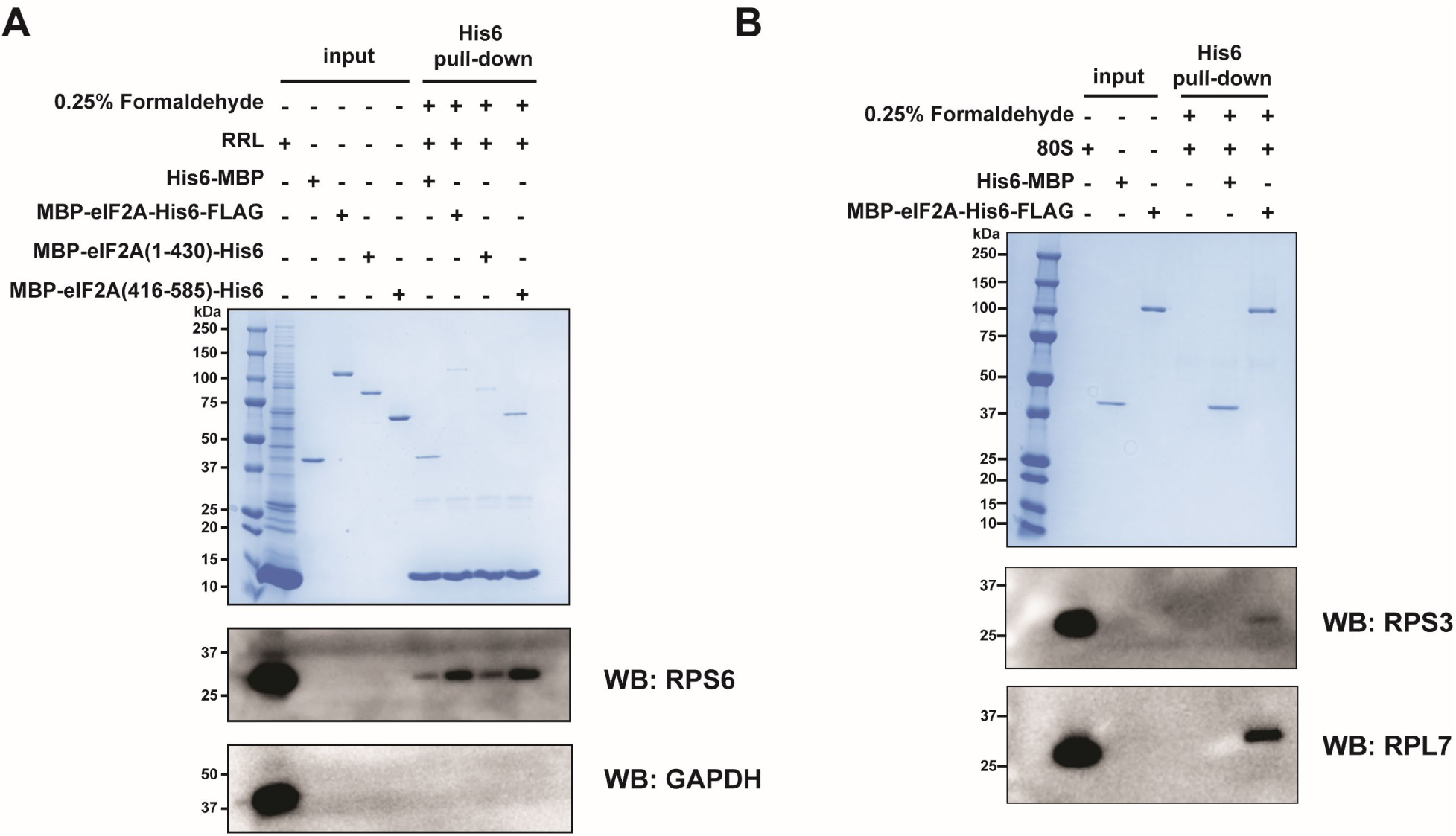
Control pulldown experiments with deletion mutants and 80S ribosomes. A) SDS-PAGE, Coomassie stain, and Western blot analysis to assess the ability of recombinant His6-MBP and the indicated recombinant His6-tagged eIF2A proteins to pulldown 40S subunits from RRL with Ni^2+^-NTA magnetic beads. 0.25% (v/v) formaldehyde was included in the indicated samples. RPS6 was used as a marker for 40S subunits. GAPDH was used as a negative control. Coomassie staining was used to confirm capture of His6-tagged recombinant proteins. Unequal pulldown was noted between full-length and deletion mutants in these cross-linking conditions. B) SDS-PAGE, Coomassie stain, and Western blot analysis to assess the ability of recombinant His6-MBP or MBP-eIF2A-His6-FLAG to pulldown purified 80S ribosomes with Ni^2+^-NTA magnetic beads. 0.25% (v/v) formaldehyde was included in the indicated samples. RPS3 and RPL7 were used as markers for 40S and 60S subunits, respectively. Coomassie staining was used to confirm capture of His6-tagged recombinant proteins.

## Reporter mRNA sequences

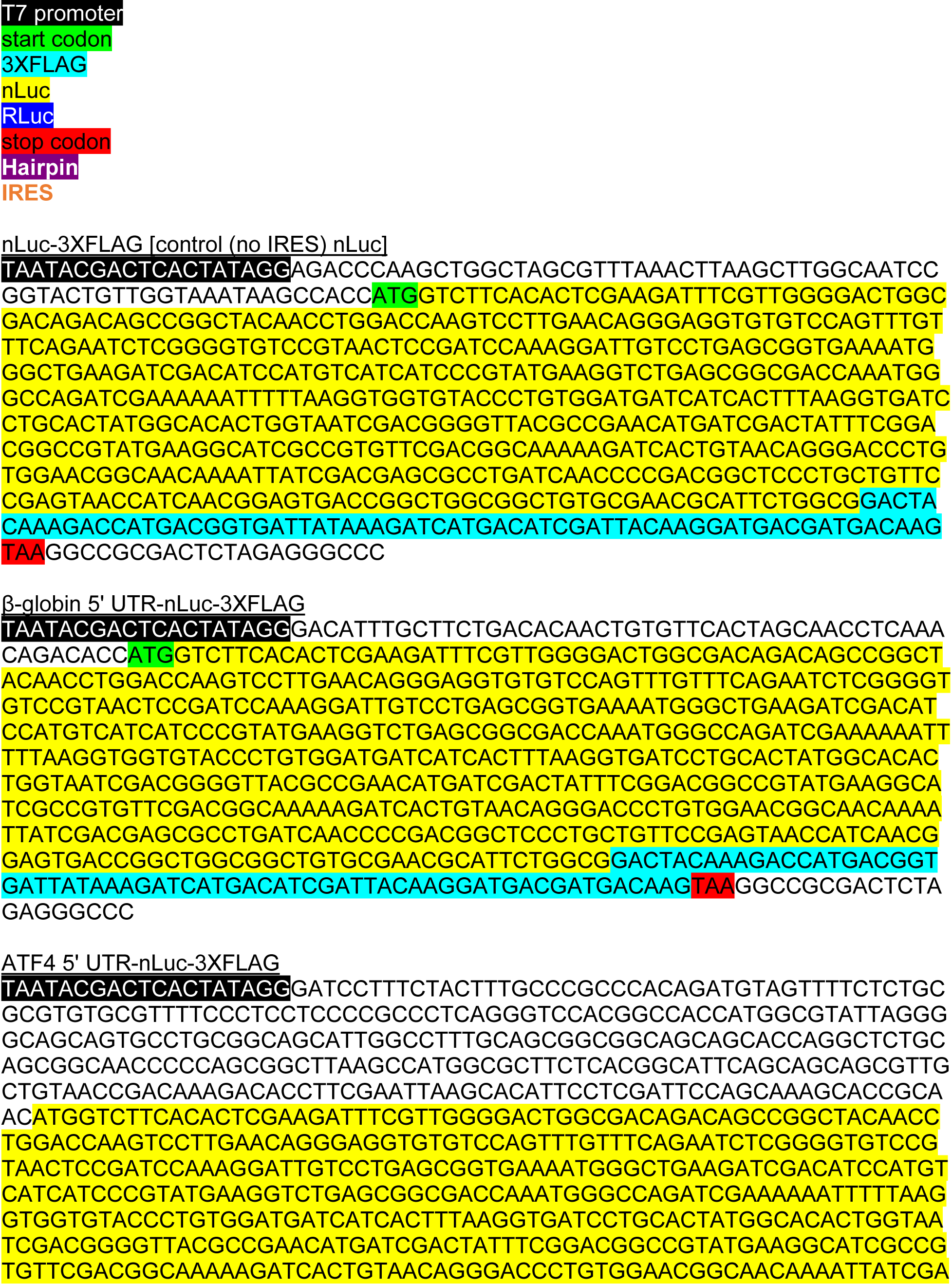
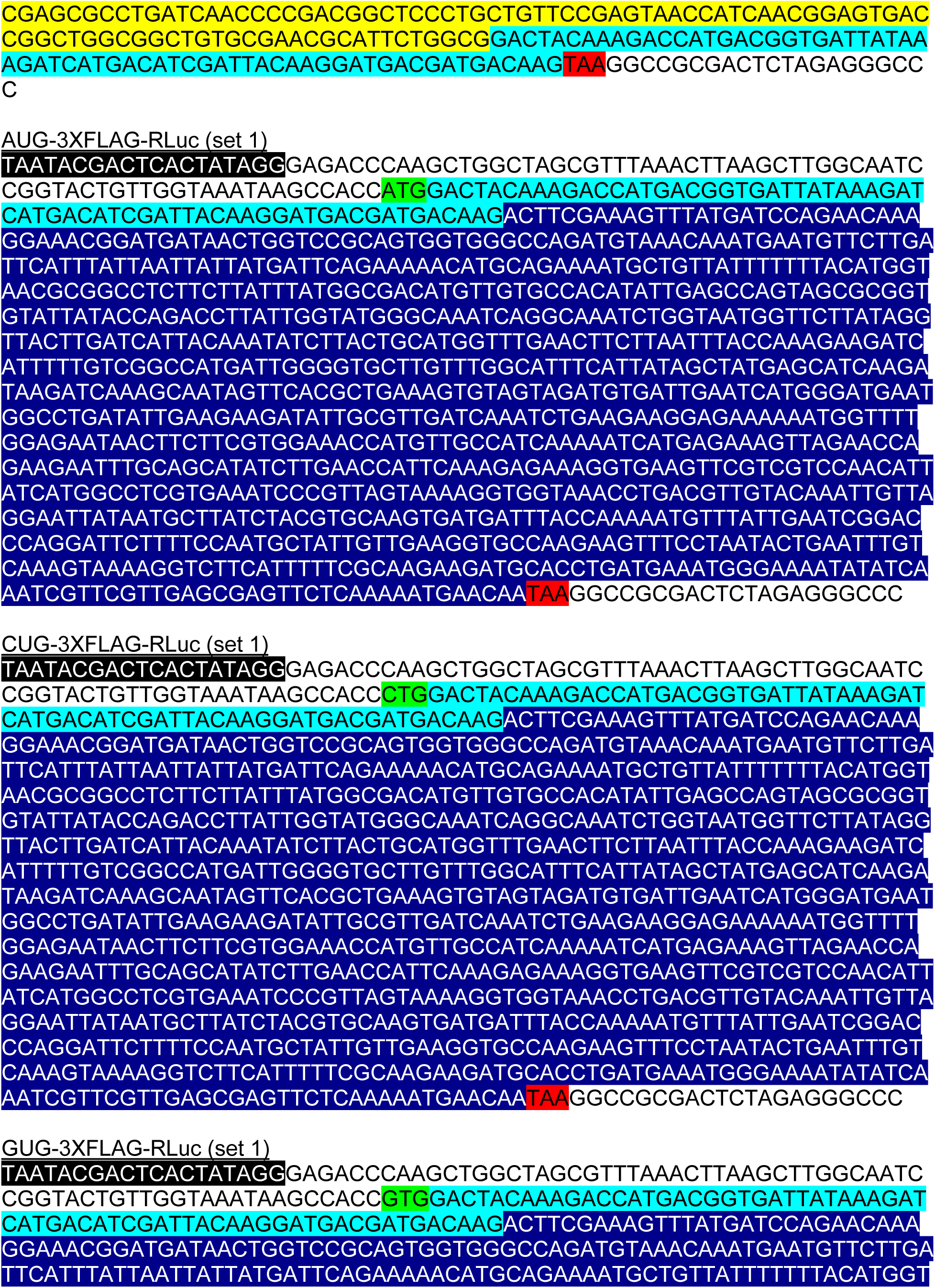
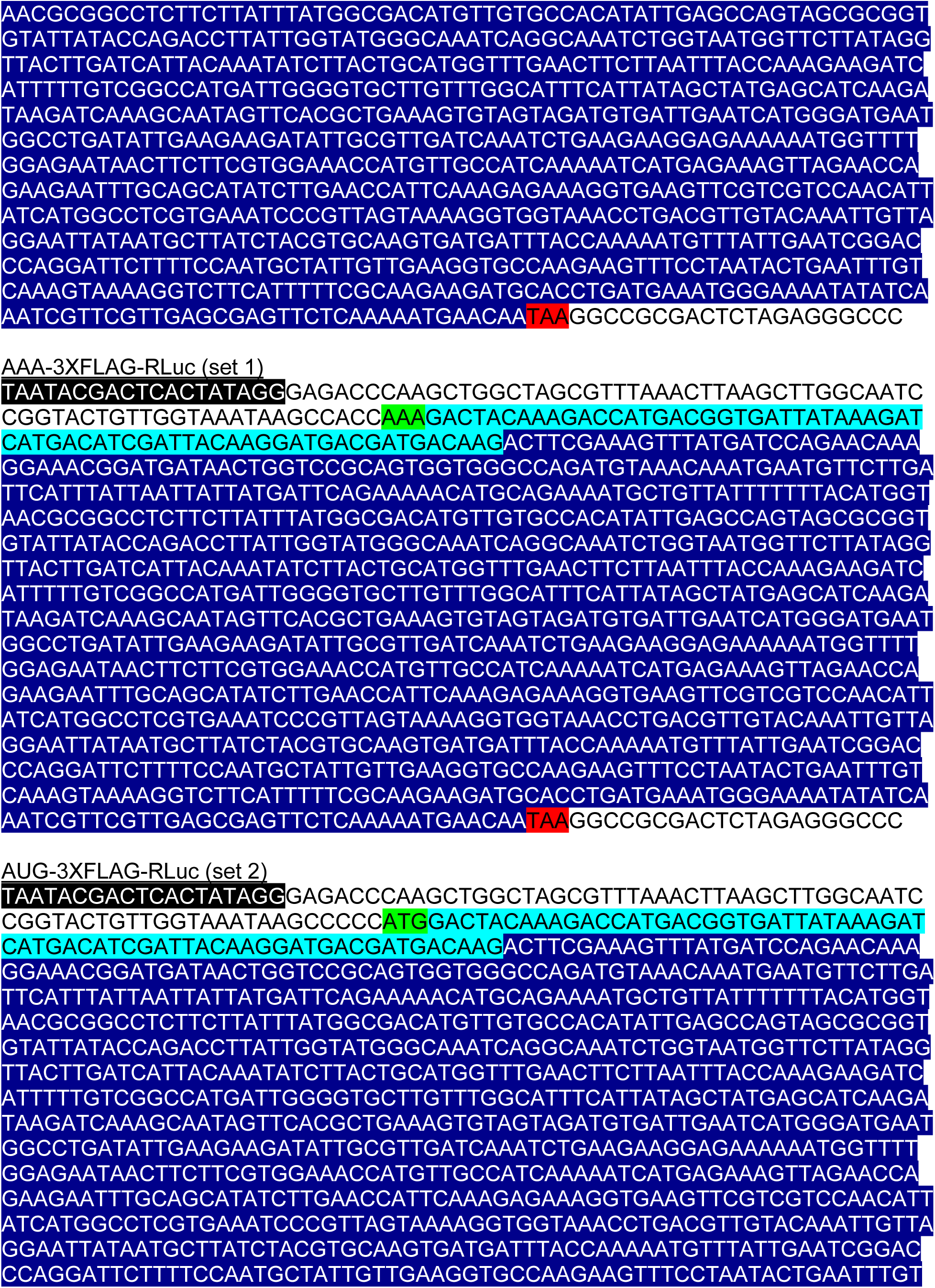
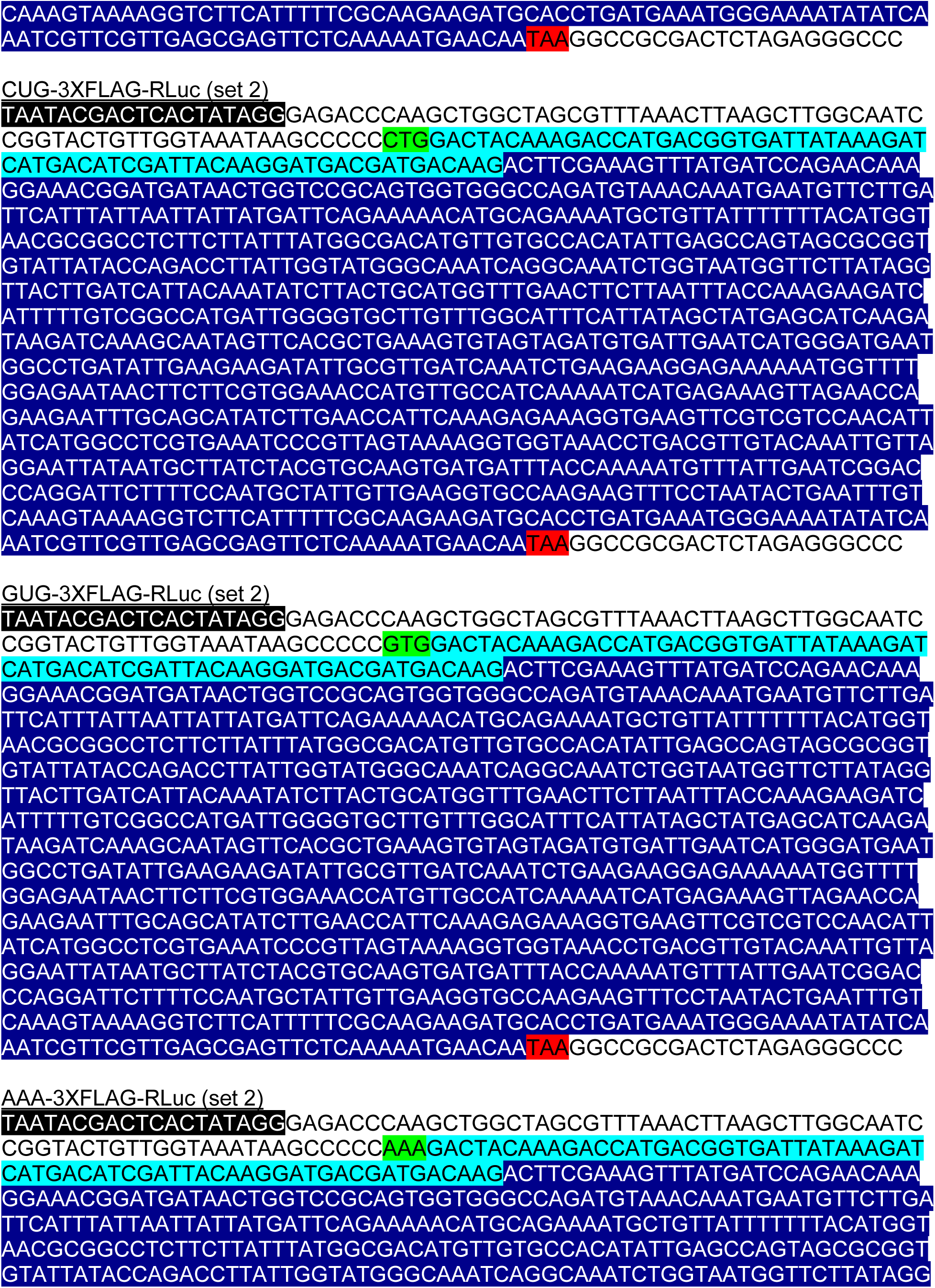
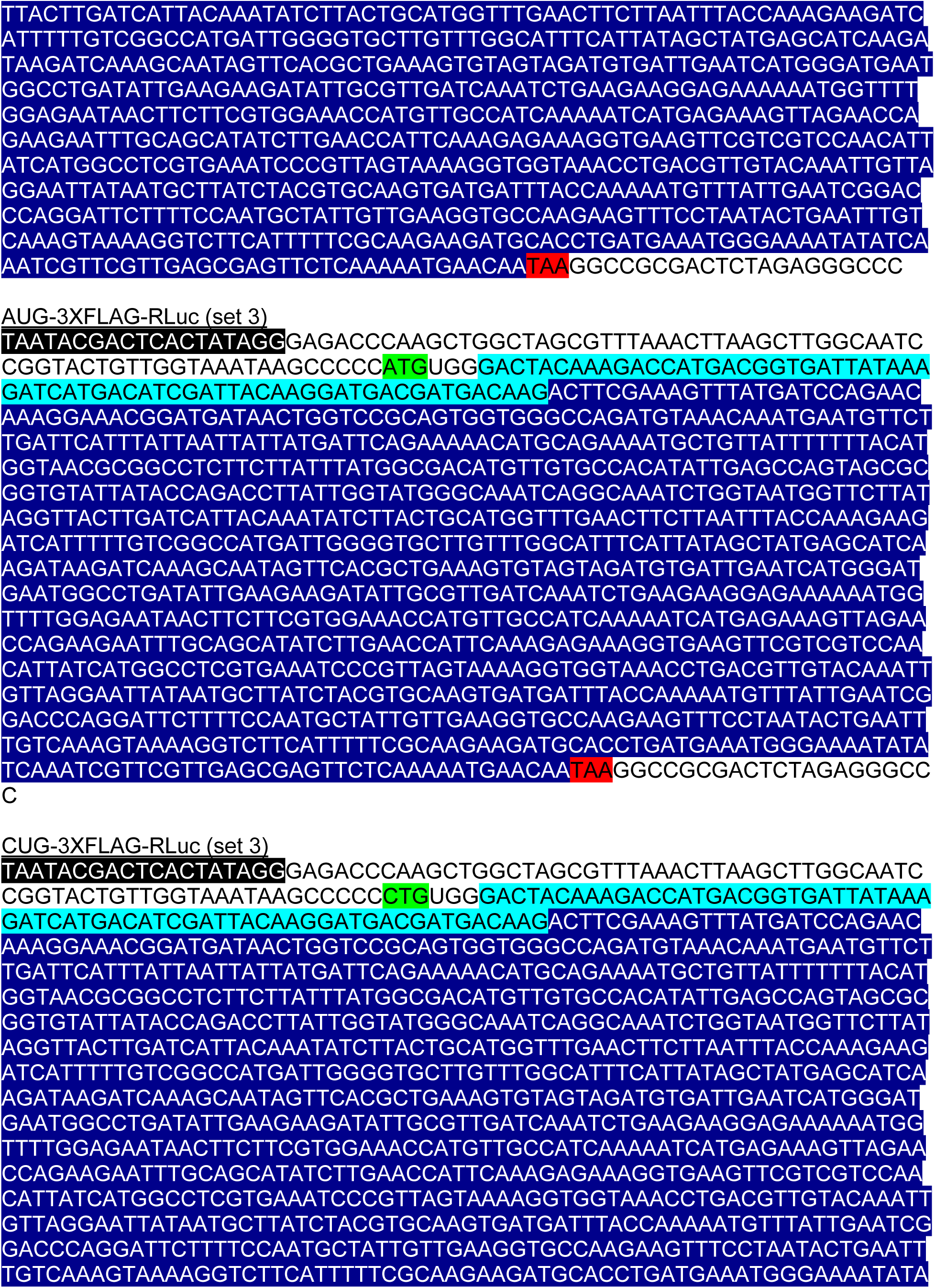
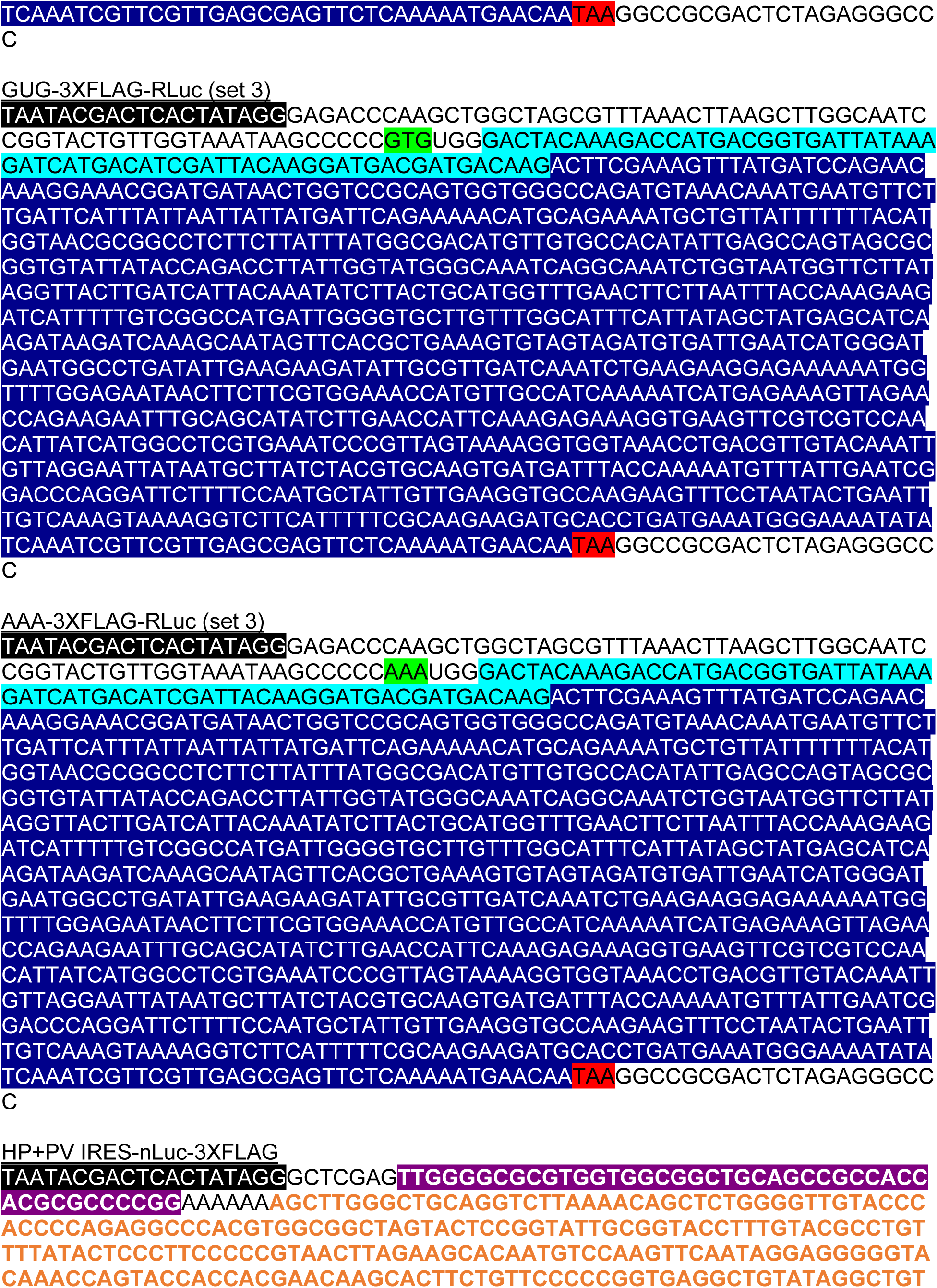
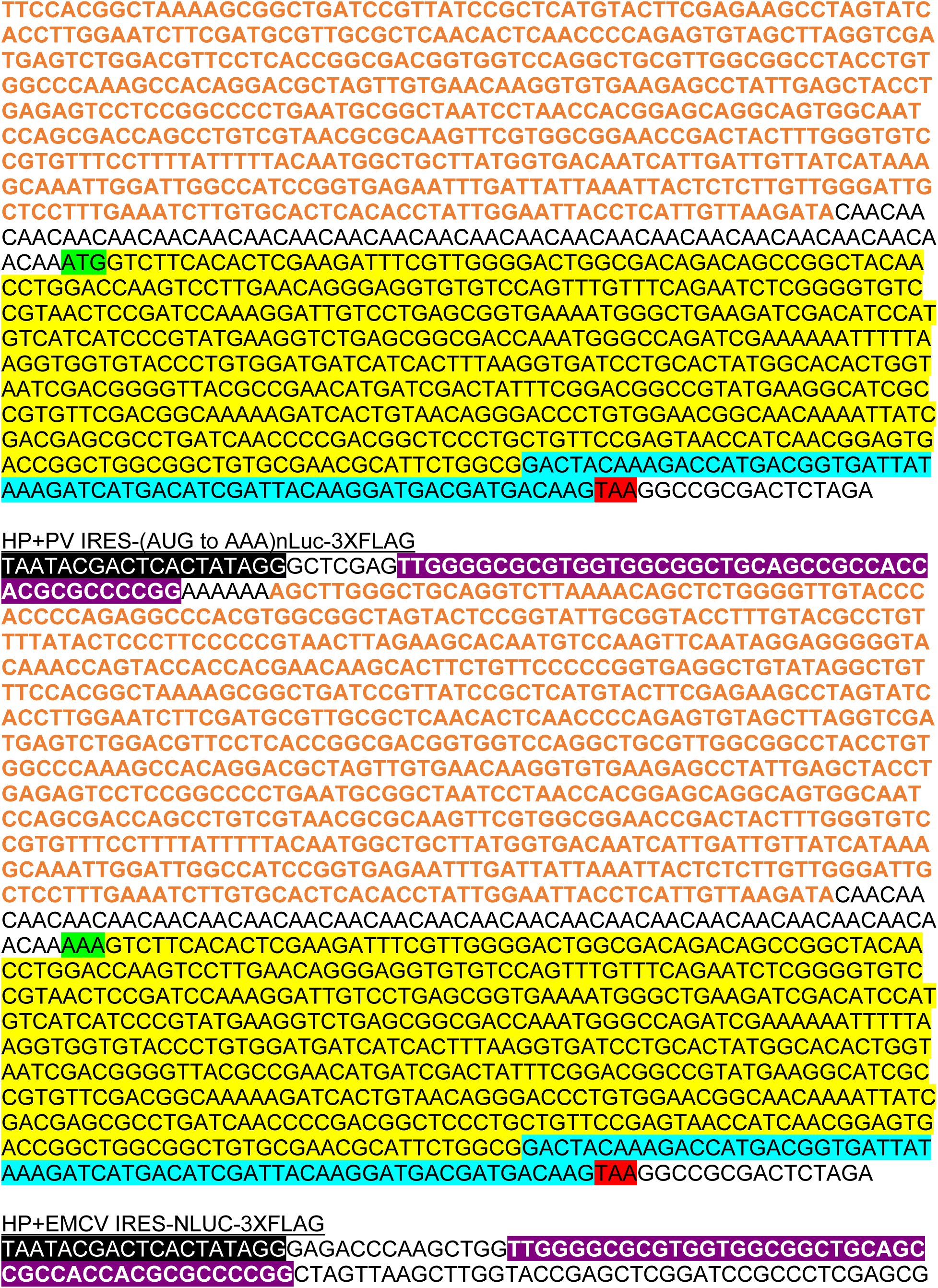
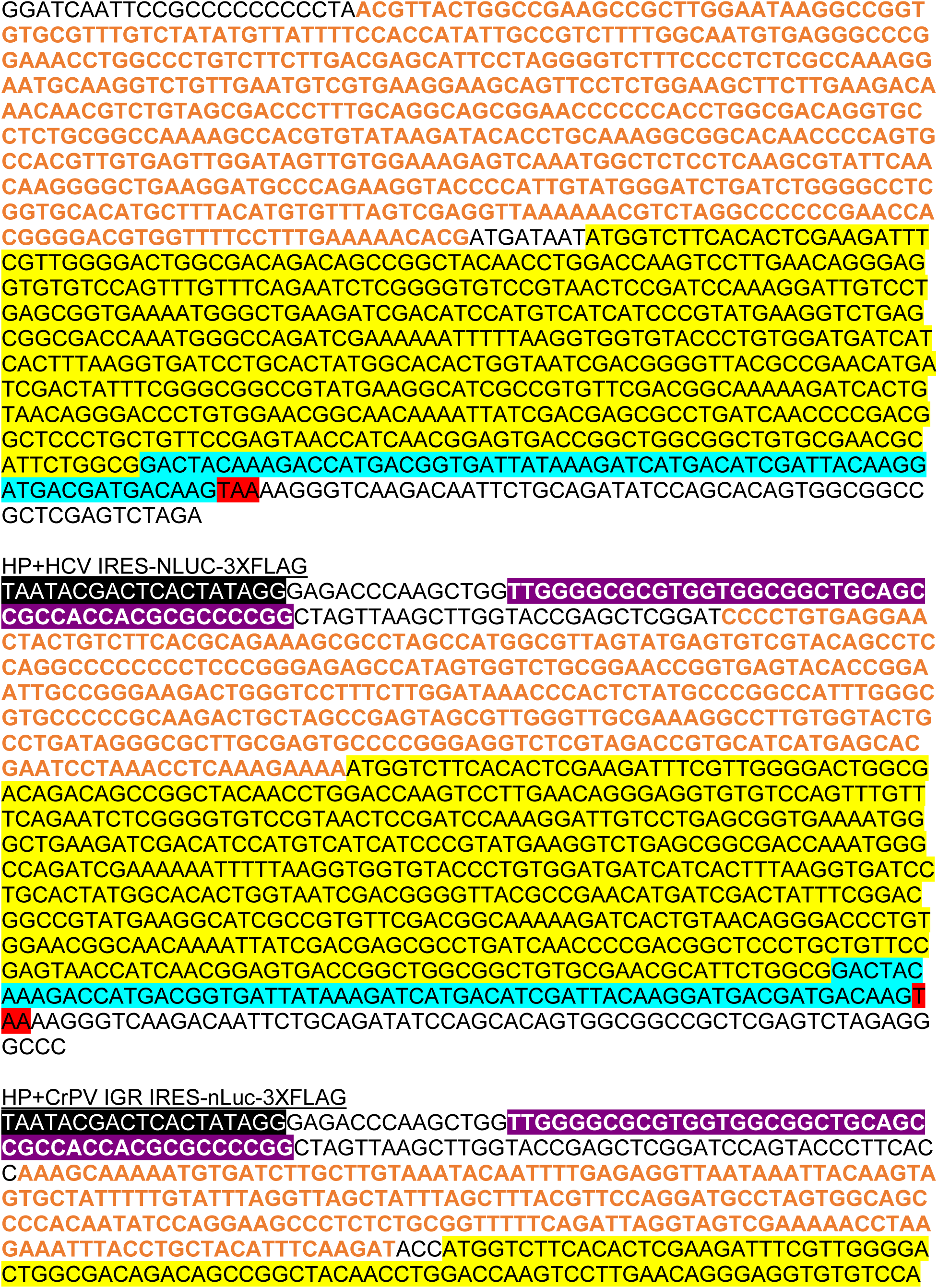
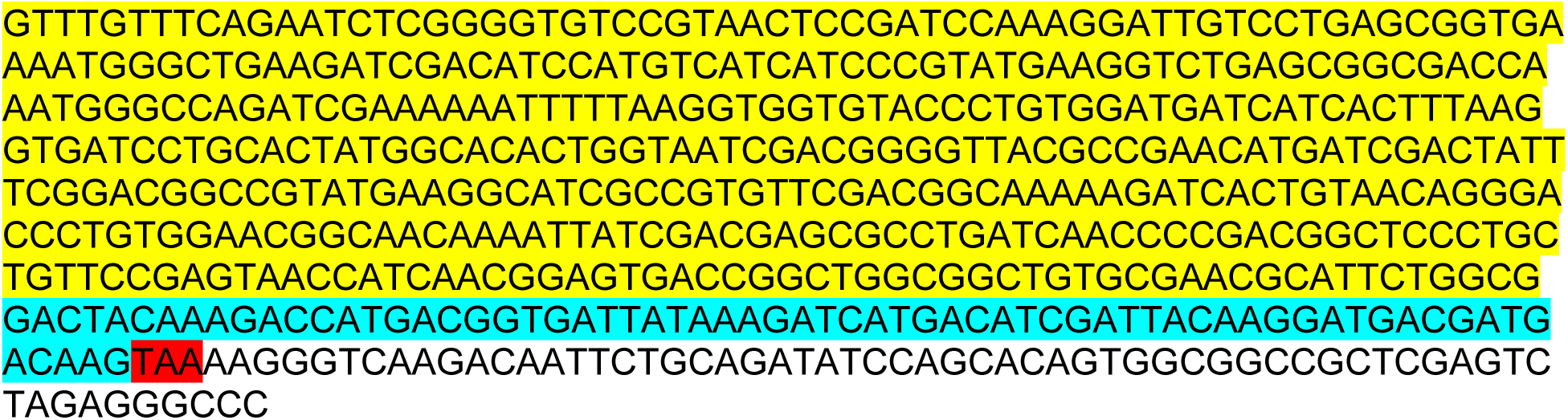

## Recombinant protein sequences

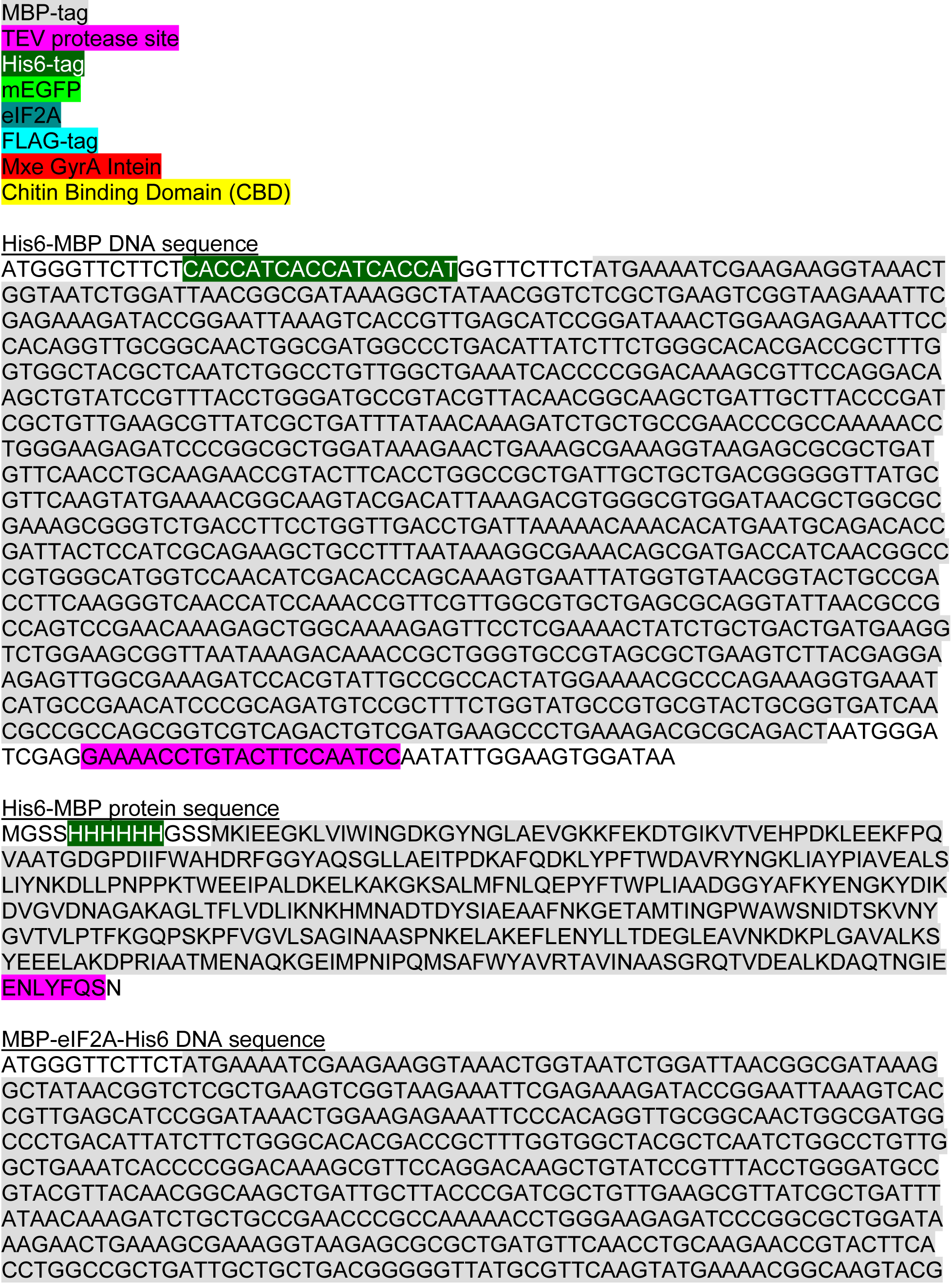
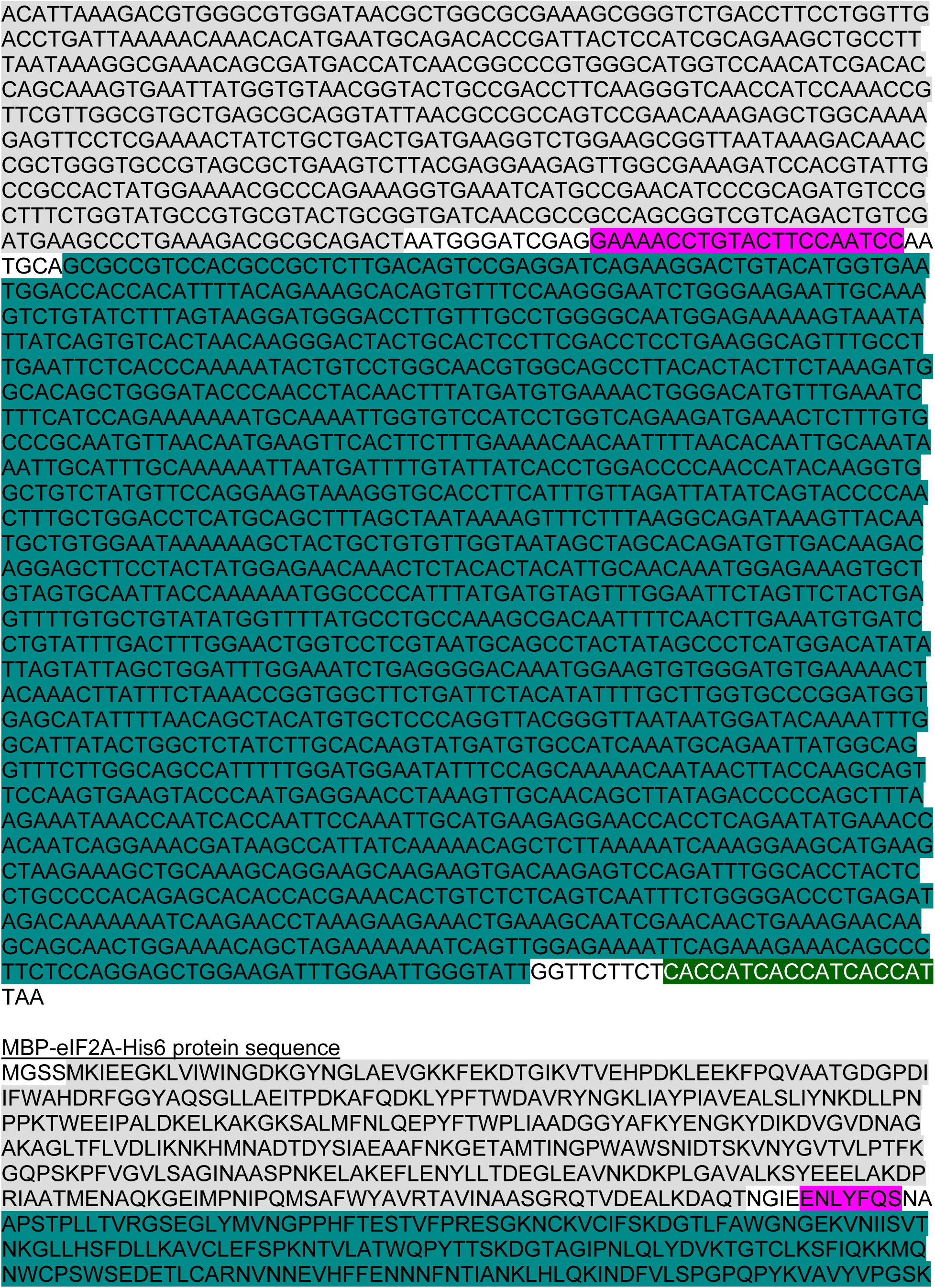
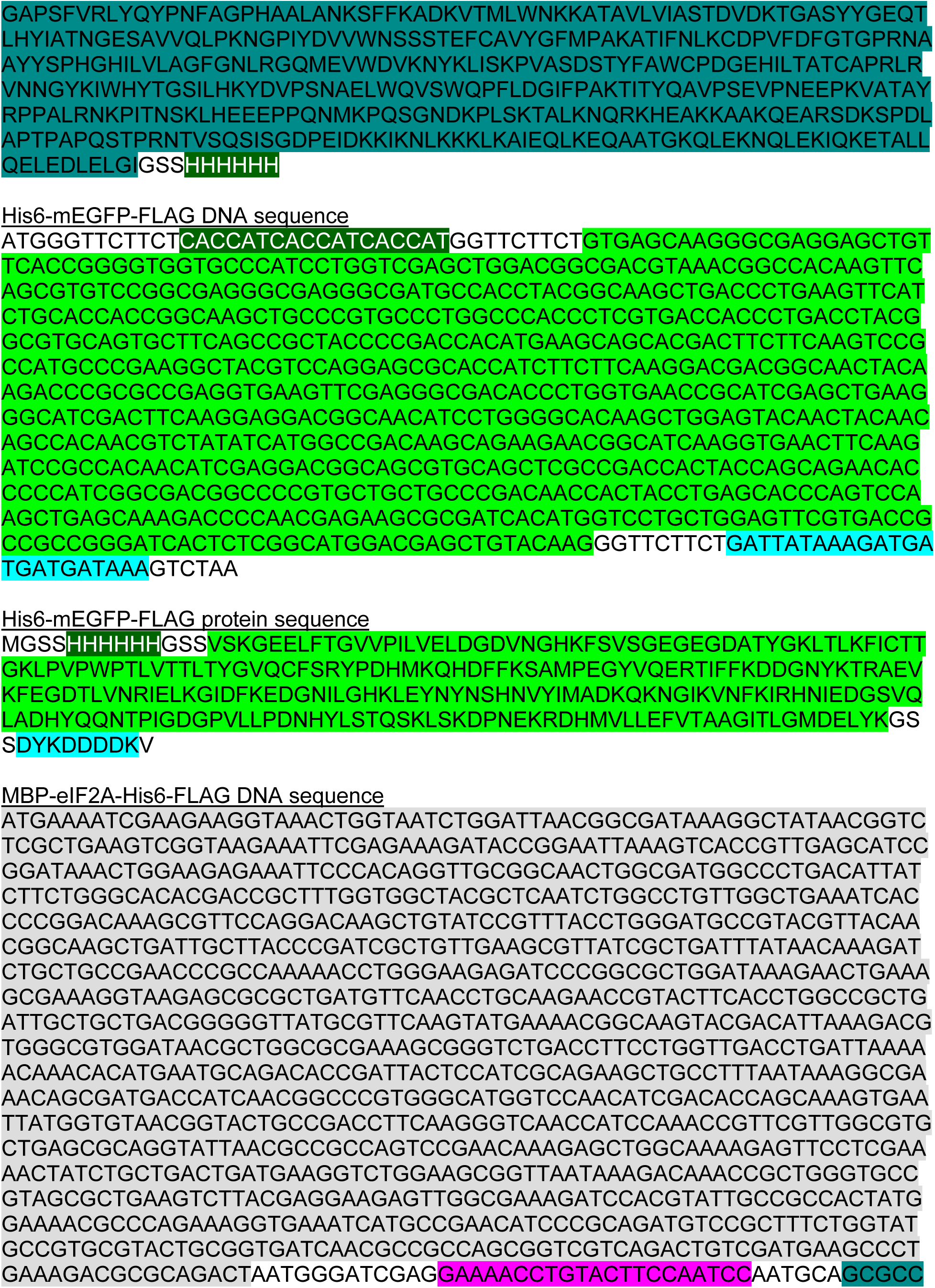
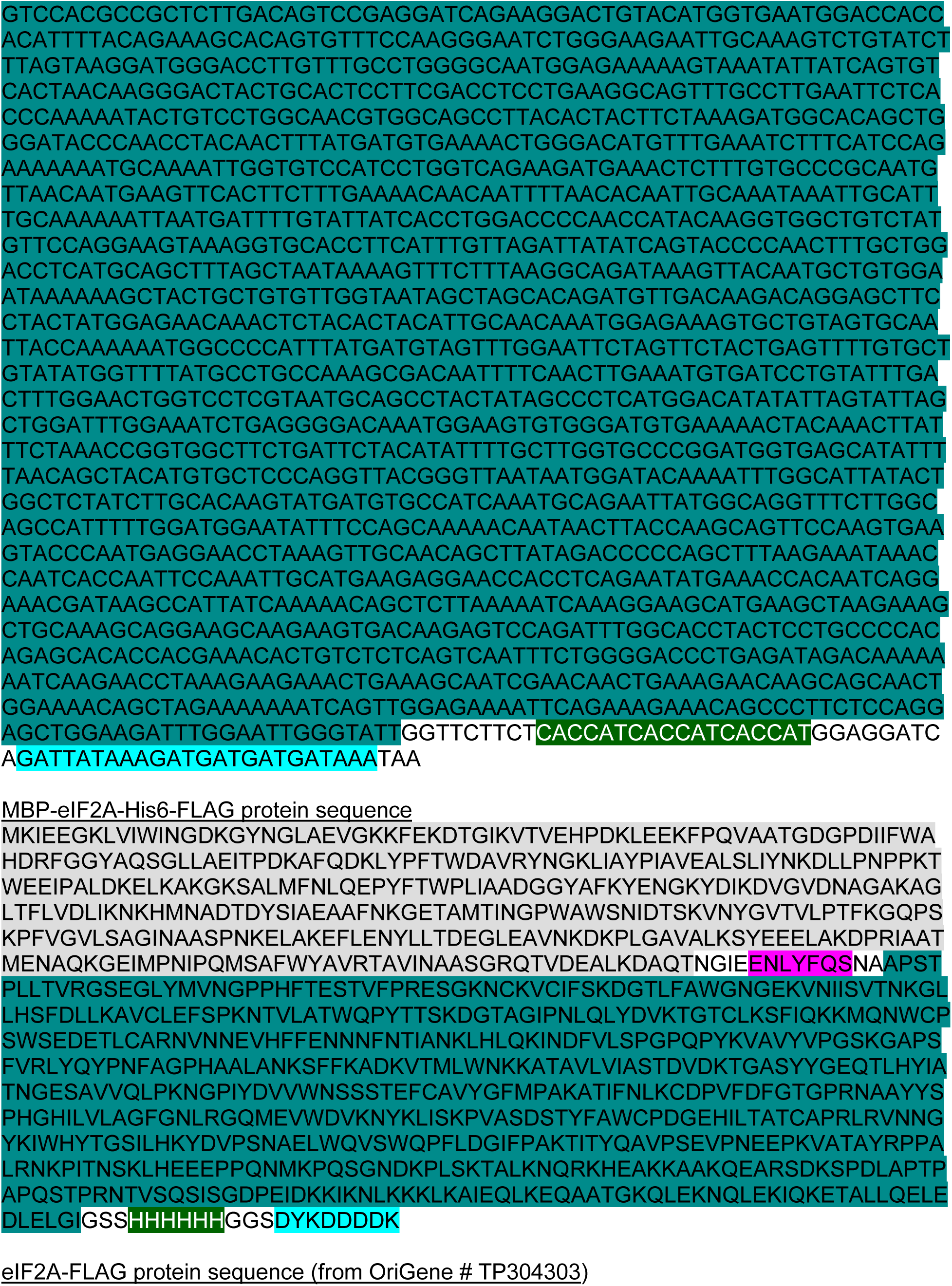
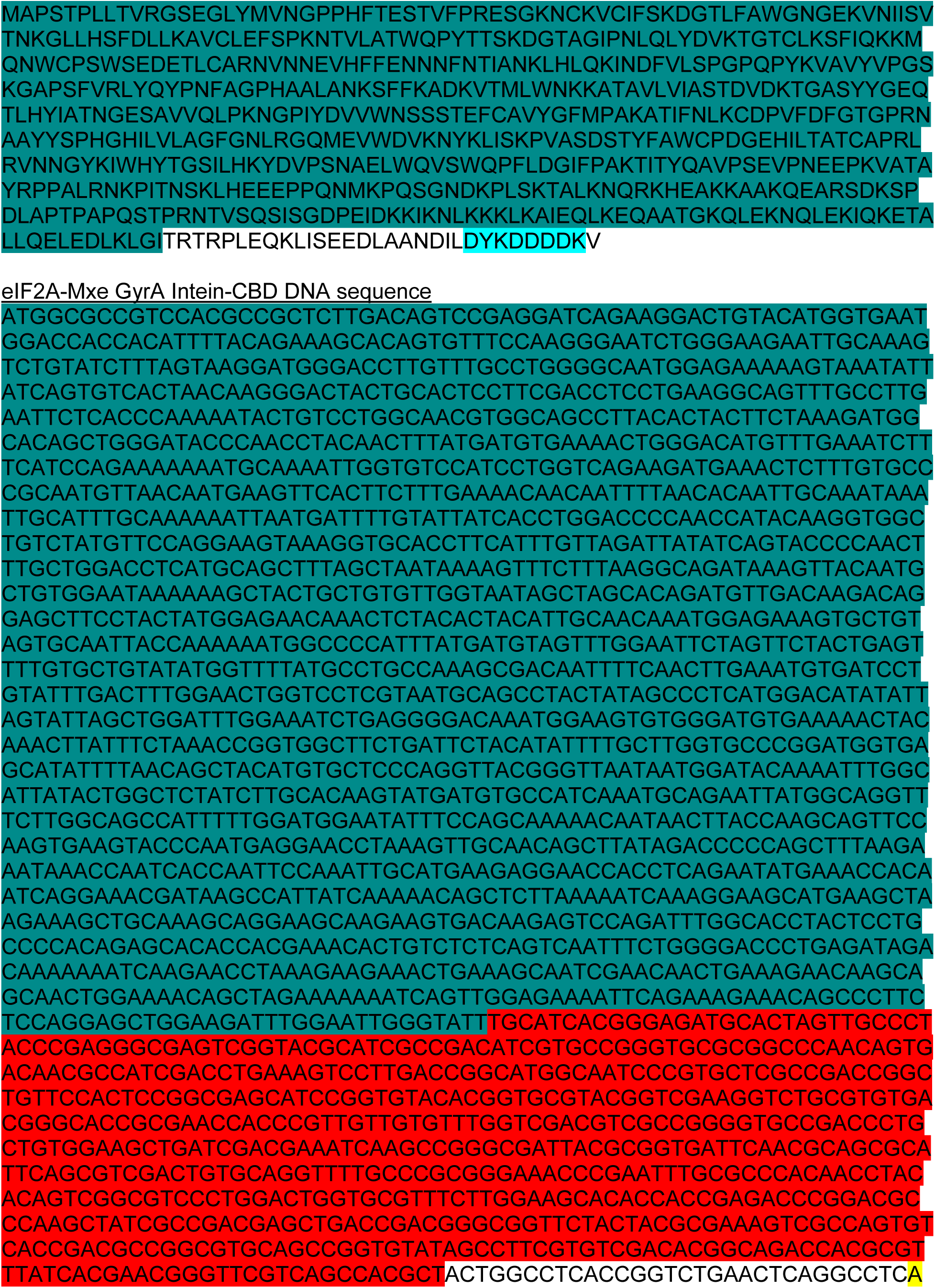
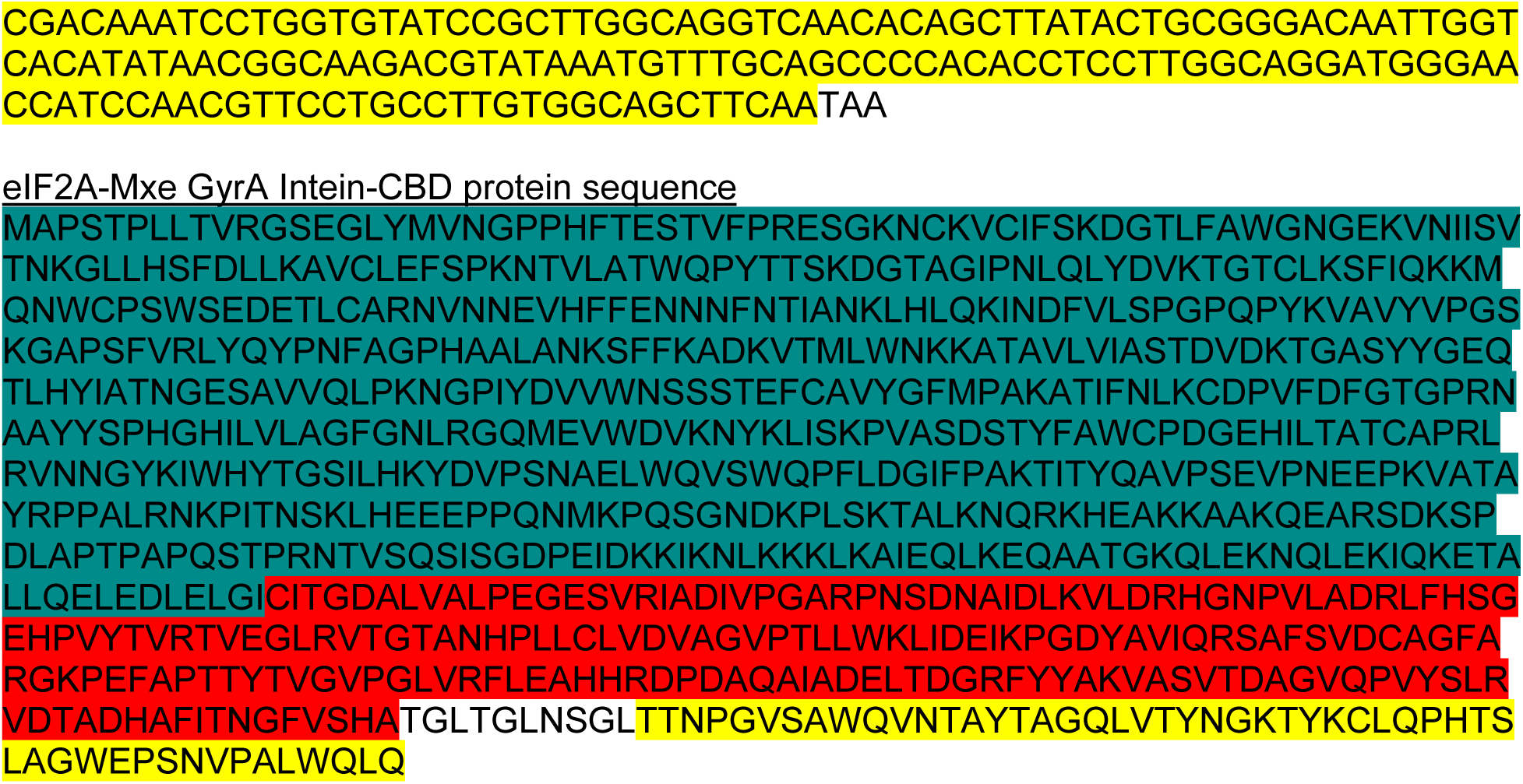

